# A genomic catalog of Earth’s bacterial and archaeal symbionts

**DOI:** 10.1101/2025.05.29.656868

**Authors:** Juan C. Villada, Yumary M. Vasquez, Gitta Szabo, Ewan Whittaker-Walker, Miguel F. Romero, Sarina Qin, Neha Varghese, Emiley A. Eloe-Fadrosh, Nikos C. Kyrpides, SymGs data consortium, Axel Visel, Tanja Woyke, Frederik Schulz

## Abstract

Microbial symbiosis drives the functional and phylogenomic diversification of life on Earth, yet remains underexplored due to culturing challenges. This study employed machine learning (ML) to predict symbiotic lifestyles in hundreds of thousands of microbial genomes from diverse environmental metagenome samples and reference genomes. Predictions were performed using `symclatron`, a novel ML framework developed here to identify genomic signatures of symbionts. Predictions were deposited in a novel Symbiont Genomes catalog (SymGs). The results indicated that 15-23% of uncultivated microbes likely engage in symbiotic relationships with other organisms and are present in half of all known bacterial and archaeal phyla. We also identified genomic signatures of symbiotic lifestyles, including the loss of certain metabolic functions and the differential presence of others that may enable host-dependent living. The `symclatron` software and the SymGs catalog represent valuable resources for studying symbioses, facilitating future mechanistic investigations of host-microbe associations.

## 2. INTRODUCTION

The short- and long-term intricate relationships between microorganisms and their hosts, known as microbial symbioses, are fundamental to the functioning of life on Earth (Douglas 2011; Human Microbiome Project Consortium 2012; Douglas 2014). These dynamic relationships span a spectrum from mutualism to parasitism (Drew, Stevens, and King 2021) and are integral to the health and evolution of all partners involved (Bordenstein and Theis 2015). On the one hand microbial symbionts play critical roles in processes such as nutrient cycling (Kipp et al. 2023; Rädecker et al. 2023), transfer of energy in the form of ATP (Graf et al. 2021), enhancement of host stress resistance (Hector et al. 2022), and disease suppression and protection from pathogens (Daisley et al. 2020; Villada and Schulz 2022; Zhao et al. 2023). On the other hand, microbial pathogens infect eukaryotic cells and adversely affect their host fitness (Salje 2021). The genetic makeup of symbionts often determines host range and interaction (Drew, Stevens, and King 2021; Perreau and Moran 2022), however, the availability of symbiont genomes is skewed towards those that are animal- or human-associated or amenable to laboratory cultivation (Relman 2011; Husnik et al. 2021; Delaux and Schornack 2021). This bias has caused researchers to overlook the true breadth of symbiont diversity in nature, potentially missing novel and globally significant symbiont clades yet to be identified.

Recent advances in metagenomics have provided the tools to capture and analyze the vast array of genetic material present in environmental samples, offering unprecedented insights into the microbial world (Koonin, Makarova, and Wolf 2021). The ability to reconstruct population genomes, so-called metagenome-assembled genomes (MAGs), and single amplified genomes (SAGs), from environmental sequencing data has revolutionized our understanding of microbial diversity, functional capabilities and ecology (Rinke et al. 2013; Brown et al. 2015; Parks et al. 2017; Castelle et al. 2018; Nayfach et al. 2021; Pavlopoulos et al. 2023), and the evolution of the eukaryotic cell (Stairs et al. 2020; Eme et al. 2023). To date, hundreds of thousands of MAGs have been generated (Parks et al. 2022; Gurbich et al. 2023; I.-M. A. Chen et al. 2023). While MAG taxonomy can be linked to their predicted metabolic traits, the lifestyles of these organisms remains largely unknown. Taxonomy alone is often insufficient to predict lifestyle due to variation within related groups and the prevalence of novel organisms in MAG datasets. Analyzing genomic features related to function and dependency offers a more direct approach (Guerrero-Egido et al. 2024). In particular genomes of symbiotic clades have been disproportionately underrepresented in large-scale MAG studies. This bias occurs because estimates of genome completeness often fall below the medium-quality threshold for these genomes (McCutcheon and Moran 2011; Bowers et al. 2017; Chklovski et al. 2023). The concept of the symbiome encompasses the entire assemblage of co-localized and co-evolving organisms, including both host and symbionts (Tripp et al. 2017). However, a linkage between symbiont and host can often not be established in complex metagenomic data sets. Yet, it is plausible that many MAGs and SAGs represent uncultivated microbial symbionts, and therefore, a symbiont-centric view of the global microbiome data could represent a powerful approach to uncover the full extent of the genomic underpinning of bacterial and archaeal symbionts of eukaryotes.

Here, we leverage machine learning models trained on large-scale genomic data to map the microbial symbionts of eukaryotes across the globe. Operationally, we distinguish potential symbionts from free-living microbes based on genomic features reflecting lifestyles ranging from “host-associated” (encompassing ectosymbionts and facultative endosymbionts) to the more dependent “intracellular” symbionts, excluding syntrophic partnerships (See Methods section “Definition of symbionts, host-associated, and obligate intracellular”). We investigate the prevalence of predicted symbionts across the microbial tree of life, systematically detect the taxonomic extent of symbiont-exclusive clades, and analyze the functional and metabolic patterns that characterize host-associated and intracellular lifestyles. The results of our large-scale approach enrich our understanding of microbial symbiont prevalence, diversity and metabolic capacity, providing unprecedented global insights into microbial symbiosis.

## 3. RESULTS

### 3.1. An AI/ML framework (symclatron) for accurately classifying symbiotic lifestyles

#### 3.1.1. Symbiont genome dataset and feature extraction

We constructed a comprehensive symbiont proteome database and extracted genomic features to enable lifestyle classification (Figure 1). The first part included a reference set of 792 symbiont proteomes from across major clades (Figure 1A) that was used to capture functional signals of symbiosis. We performed orthogroup inference on these symbiont proteomes, from which we built 20,063 high-confidence profile hidden markov models (HMMs) for orthogroups that contained ≥5 member proteins. The HMMs serve as potential markers of symbiotic gene content (Figure 1A). Next, we built a dataset comprising 6,751 labeled microbial genomes (Figure 1B) available in the IMG/M database (I.-M. A. Chen et al. 2023), spanning three lifestyle classes: free-living (*n* = 5,959), host-associated (*n* = 409), and obligately intracellular (*n* = 383). See detailed definitions of these lifestyle labels in the subsection “Definition of symbionts, host-associated, and obligate intracellular” of Methods. With the aim of accounting for the effect of genomes incompleteness in the downstream training of our models, all the genomes were artificially reduced to different levels of genome completeness and fragmentation lengths (Figure 1B). We then applied uniform genome gene calling and annotation across the entire labeled dataset, yielding ∼244 million proteins (Figure 1B). Each protein was scanned against the symbiont HMM library (Figure 1A) to determine orthogroup memberships. As a result, each genome was represented as a 20,063-dimensional feature vector reflecting the bitscore of symbiont-related orthogroups.

**Figure 1.**
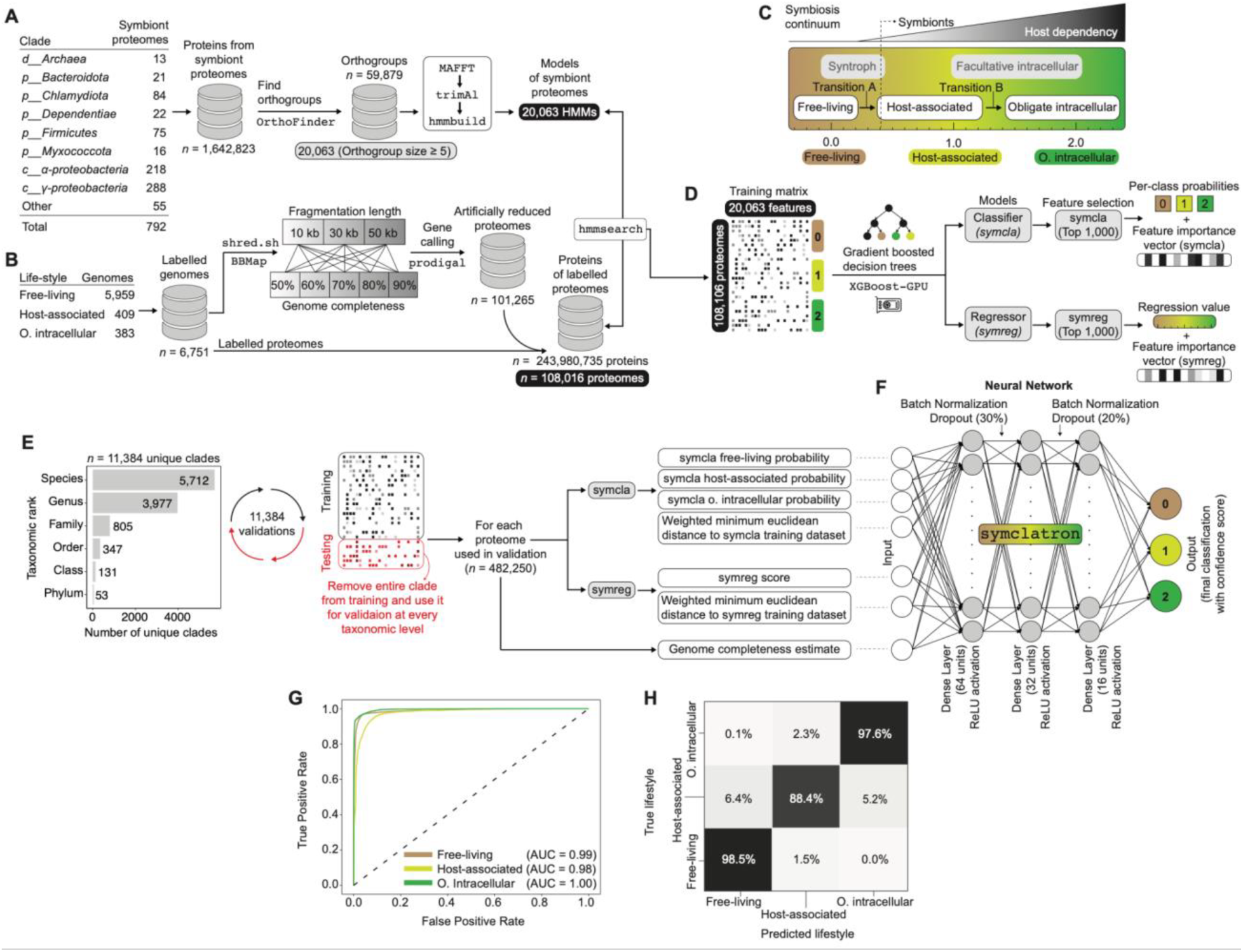
Developing an AI/ML framework for symbiotic lifestyle prediction in microbial genomes. (A) Feature engineering workflow for detecting the genomic features specific to symbiont genomes. (B) Construction of a genus-level representative database of lifestyle-labeled genomes accounting for artificial genome reductions. (C) Schematic of the symbiosis continuum concept applied in the labeling strategy for models training. (D) Construction of the training matrix for the symbiont classifier (*symcla*) and symbiont regressor (*symreg*) via extreme gradient boosted decision trees (XGBoost). The final models were optimized for the most relevant 1,000 features. (E) Strategy implemented to calibrate the final lifestyle prediction by training a neural network the output of orthogonal validations. The neural network was trained on 11,384 rounds of benchmarking and validations of the classifier and regressor models by blinding the models to entire clades from the training data. The barplot on the left shows the total number of unique clades analyzed at each taxonomic rank. (F) Architecture of the neural network featuring batch normalization and dropout layers (30% and 20%), with three dense layers using ReLU activation. The output layer also yields confidence scores for each lifestyle class (0 = free-living, 1 = host-associated, 2 = obligate intracellular). (G) Receiver Operating Characteristic (ROC) curves showing classification performance for each lifestyle category (one-vs-rest). (H) Confusion matrix at 0.725 confidence threshold depicting correct classifications for each category. The data split for the neural network validation was done at 80% for training (*n* = 385,800 proteomes) and 20% for testing (*n* = 96,450 proteomes).

Genome labelling was done following our conceptual demarcation of the symbiosis continuum (Figure 1C). Then, the 20,063-dimensional encoding captures each genome’s potential host-dependence traits (Figure 1C). Accordingly, we used the encodings to train a symbiont classifier model (`symcla`) based on discrete class labels, and to train a regressor model (`symreg`) derived from the continuous labels (Figure 1D). To optimize computational efficiency, the top 1,000 most relevant features were selected for each model.

#### 3.1.2. Calibrating predictions with clade-level cross-validations and the neural network symclatron

To benchmark the model’s generalizability to novel taxa, we employed a clade-level leave-one-clade-out cross-validation strategy (Figure 1E). In total, 11,384 unique clades (across species, genus, family, order, class, and phylum levels) were represented in the labeled dataset (Figure 1E, left). For each validation run, we withheld all genomes from one clade (e.g., all members of a given family) from training and then tested the model on those held-out genomes (Figure 1E, right). This process was repeated such that each clade (at each taxonomic rank) was left out in turn, yielding 11,384 clade-specific validations. By removing entire lineages during training, this rigorous scheme evaluates how well the classifier can predict lifestyles for previously unseen lineages.

We developed a feed-forward neural network, termed “`symclatron`”, to provide a final prediction of each genome’s lifestyle based on the clade-level cross-validation data and other interpretable genomic features (Figure 1F). Rather than using thousands of raw ortholog features directly, the `symclatron` model integrates seven informative features per genome that summarize its position in symbiotic gene-content space. These features capture both the categorical and continuous signals of symbiosis, as well as genome quality, and are defined as follows (Figure 1F):

`symcla` class probabilities (3 features): The probabilities of the genome being free-living, host-associated, or obligate intracellular, as predicted by an initial gradient-boosted trees classifier trained on the 20,063-dimensional feature matrix.

`symreg` score (1 feature): A continuous symbiosis score ranging from 0.0 (free-living) to 2 (obligate intracellular) produced by a gradient-boosted regression model (`symreg`). This models the genome’s position along the host-dependence continuum (Figure 1C).

Distance to training datasets (2 features): For both the `symcla` classifier feature space and the `symreg` feature space, we computed the genome’s weighted Euclidean distance to the nearest training-set genome. These distance metrics (weighted by feature importances from the symcla/symreg models) quantify how atypical or novel the genome’s gene content is relative to known symbionts and free-living organisms.

Genome completeness (1 feature): An estimate of genome completeness (assembly quality) for the proteome, included to ensure that partial genomes are appropriately handled.

All seven features were scaled and passed as input to the neural network. The `symclatron` architecture consists of fully connected dense layers with batch normalization and dropout regularization (Figure 1F). Specifically, two hidden layers (each with ReLU activation and dropout rates of 30% and 20%, respectively) transform the input features, followed by an output layer with a softmax activation that yields probability scores for each of the three lifestyle classes (0 = free-living, 1 = host-associated, 2 = obligate intracellular). This design leverages biologically interpretable inputs, derived from gene content patterns and host-dependence signals, while using the neural network to capture non-linear combinations of these inputs for final prediction. By building on the intermediate `symcla` and `symreg` models (which themselves provide feature importance insights), the `symclatron` predictions remain grounded in tangible biological features rather than opaque latent variables.

#### 3.1.3. Classification performance across lifestyles

The `symclatron` model achieved high classification accuracy for all three lifestyle categories in cross-validation (Figure 1G-H). Receiver operating characteristic (ROC) analysis (Figure 1G) showed that the model discriminates lifestyles with excellent performance: the one-vs-all AUC was 0.99 for free-living, 0.98 for host-associated, and 1.00 for obligate intracellular genomes. These high AUC values indicate that `symclatron` consistently ranks the true class higher than the alternatives across diverse genomes. Correspondingly, the classifier’s confusion matrix confirms accurate assignment of lifestyles (Figure 1H). Overall accuracy was 96%, with most genomes falling on the diagonal of the confusion matrix. Free-living bacteria were almost always correctly identified (98.5% of free-living genomes predicted as free-living, Figure 1H), with only minimal misclassification (1.5% predicted as host-associated and effectively none as intracellular). Obligate intracellular symbionts were also recognized with high precision (97.6% correctly classified; Figure 1H). The model exhibited a slight tendency to confuse the two symbiotic classes in a minority of cases: about 5.2% of host-associated genomes were mis-predicted as obligately intracellular, and 2.3% of obligate intracellular genomes were predicted as host-associated (Figure 1H). This minor class mixing reflects the biological reality that some host-associated bacteria approach an intracellular lifestyle, but importantly very few symbionts were mistaken for free-living organisms, or vice versa (Figure 1H). The strong performance on held-out clades underscores that the thoroughly calibrated `symclatron` can generalize to novel taxa, even when evaluating genomes from entirely untrained lineages, the model maintained robust accuracy with only rare errors. Our results demonstrate that the neural network, informed by independent models built on gene-content features, can reliably distinguish free-living vs. symbiotic lifestyles and even delineate the degree of host dependence among symbionts.

#### 3.1.4. Prediction confidence

To translate `symclatron`’s predictions into actionable biological insights, we calibrated the model’s probability outputs to assess prediction confidence (Supplemental Figure S1). We first examined the distribution of prediction confidence scores (the highest class probability for each prediction) for correct vs. incorrect assignments (Supplemental Figure S1A). This analysis revealed a clear separation: the vast majority of true-positive predictions were made with high confidence, whereas false predictions tended to have lower confidence scores (usually < 0.6). Based on this distribution, we identified an optimal confidence threshold of 0.725, above which misclassifications are exceedingly rare (Supplemental Figure S1A). At this cutoff, the false-positive rate is only 1.5% (i.e. only 1.5% of incorrect predictions have a confidence ≥0.725), indicating that a prediction with ≥72.5% confidence is highly likely to be correct. Users of `symclatron` can thus choose to accept only predictions above this probability to minimize errors. Applying a confidence threshold allows a tunable trade-off between accuracy and data coverage (Supplemental Figure S1B). As the minimum required probability is raised, the model’s accuracy on retained predictions increases, while the fraction of genomes that meet the threshold (coverage) decreases.

Finally, we assessed model calibration across the full range of confidence scores by stratifying predictions into 40 bins and computing performance within each bin (Supplemental Figure S1C). This analysis showed that `symclatron`’s reported probabilities are well-aligned with actual outcomes. Higher confidence scores yield higher precision in lifestyle assignment, as reflected in the confusion matrices for each bin (Supplemental Figure S1C). In the lowest confidence bins (e.g. <0.5), the model’s predictions include a mix of errors, especially for the difficult host-associated category. In contrast, in the highest confidence bin (≥0.95 probability), predictions are almost always correct: free-living and obligate intracellular calls were >99% accurate, and even host-associated calls were ∼92% accurate. Even at moderately high confidence levels (for instance, ≥0.70), the precision for free-living and intracellular predictions already exceeds 97-98%, with host-associated predictions around ∼88% accurate (Supplemental Figure S1C). These results confirm that the model’s confidence scores are meaningful measures of confidence for practical applications of symclatron.

#### 3.1.5. symclatron availability

Our symbionts prediction framework uses biologically interpretable features of genomes to accurately predict microbial lifestyle with high confidence (Fig 1G-H, and Supplemental Figure S1C). The model is robust to taxonomic novelty (validated across thousands of clades) and provides probabilistic predictions that are well-calibrated, enabling flexible, threshold-tuned deployment. These characteristics make `symclatron` a powerful tool for characterizing the lifestyle of microbes, even in the face of incomplete genomes or previously unseen lineages, while retaining transparency and control over prediction confidence. `symclatron` is publicly available to users at https://github.com/NeLLi-team/symclatron.

### 3.2. SymGs, a global-scale genomes catalog of microbial symbionts

#### 3.2.1. Genome catalog construction and symbiont classification

We started by binning a comprehensive set of 31,152 public environmental sequencing data projects from the IMG/M database to construct a novel catalog of metagenome-assembled genomes (MAGs) referred to as “NeLLi2023”. We also collected reference genomes from IMG/M and GTDB. After QA/QC and de-replication at 95% ANI, we obtained a genome catalog encompassing 107,067 genomes of Bacteria and Archaea (Figure 2A). To predict symbiotic lineages within our catalog, we applied `symclatron`. Using the stringent confidence threshold 0.725, we identified 14,070 high-confidence symbiont genomes in the catalog, equating to 14% of all genomes in the catalog; we define the collection of high-confidence symbiont genomes as the Symbiont Genomes Catalog (SymGs) (Figure 2B). In other words, roughly one in seven genomes in the assembled dataset was predicted to represent a symbiont (Figure 2C). The symbiont classification rate was even higher when computed only for the MAGs fraction of the catalog, ranging from 15% up to 23% symbionts (Figure 2C), roughly one in five MAGs is predicted to be a symbiont.

**Figure 2.**
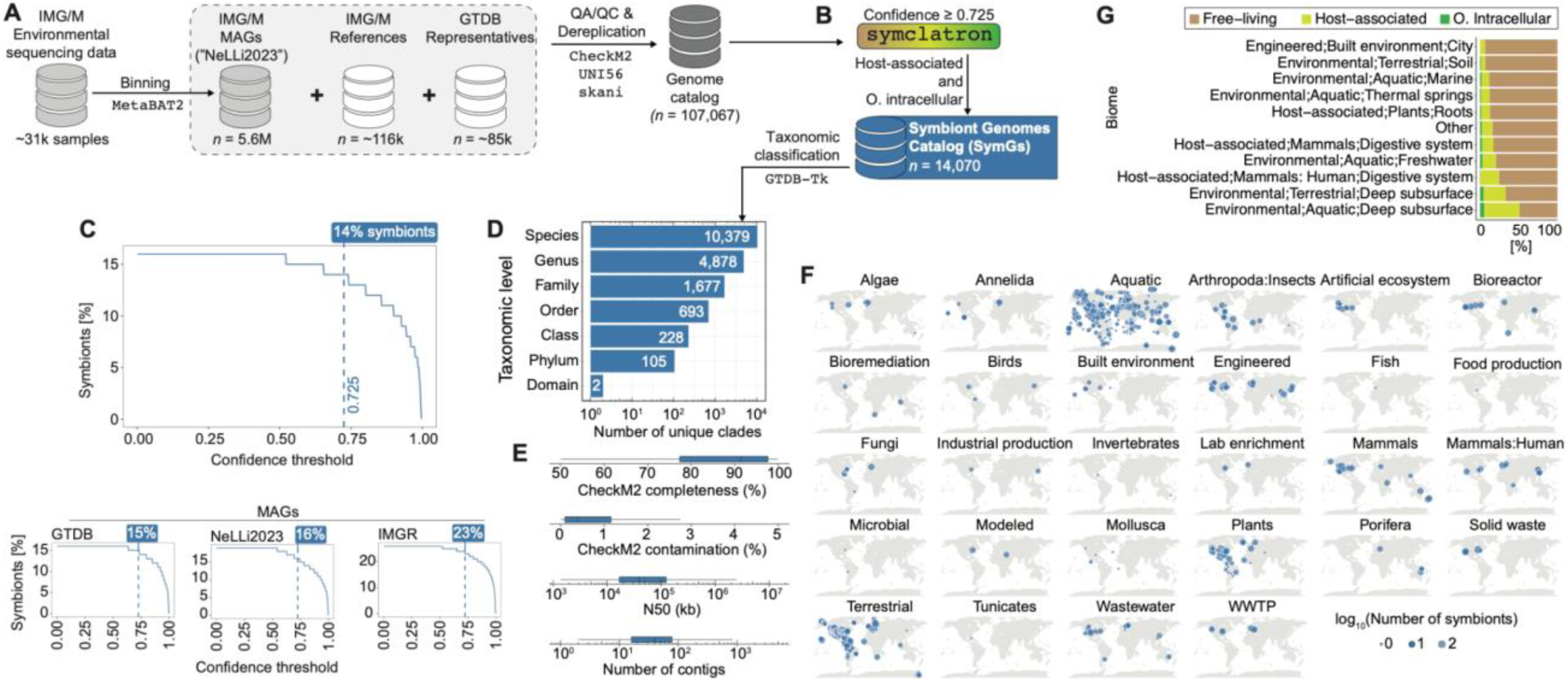
Cataloging the taxonomic and global environmental distribution of microbial symbionts. (A) Overview of the pipeline for creating the genome catalog. (B) The genome catalog was subjected to `symclatron` to predict lifestyles of Bacteria and Archaea, only high confidence predictions are reported in all downstream results. (C) Percentage of predicted symbionts as the added percentage of the categories Host-associated and Obligate intracellular across varying confidence thresholds. The bottom panels specify the percentages only for the 3 databases of MAGs used in this study. (D) Taxonomic breakdown of the clades identified in the SymGs catalog. (E) Quality assessment metrics of the SymGs catalog. (F) Global distribution of genomes in the SymGs catalog across different biomes. Geographical locations and environment data is only shown for metagenomes with available metadata in the IMG/M. (G) Percentage of predicted lifestyles (only MAGs) across the top 10 biomes with most symbiont prevalence. Remaining biomes were collapsed into the “Other” category. Bars show the relative frequency of each category in that biome.

#### 3.2.2. Taxonomic diversity and genome quality of SymGs

The catalog with 14,070 symbiont genomes (SymGs) span a wide phylogenetic range across the prokaryotic tree of life, highlighting the broad taxonomic diversity of microbial symbionts (Figure 2D). SymGs encompassed representatives from all prokaryotic ranks, including hundreds of bacterial phyla (Figure 2D). This diversity indicates that symbiotic lifestyles have arisen in many branches of Bacteria and Archaea. Indeed, the overall taxonomic profile of the SymGs closely mirrored the relative composition of the full 107,067-genome catalog (Supplemental Figure S2A), implying that most high-level clades contain at least some symbiotic members. As expected, lineages well known for symbiosis were prominent within SymGs (e.g. Pseudomonadota/Proteobacteria, Bacteroidota, Firmicutes), reflecting an enrichment of symbiont-rich clades in the catalog. At the same time, SymGs also included members of less-characterized phyla (including candidate phyla), underscoring that symbiotic associations are phylogenetically widespread and not limited to traditionally studied groups. In addition to their broad taxonomic spread, the SymGs are distinguished by high genome quality and characteristic features (Figure 2E). Most symbiont genomes were high-quality MAGs or nearly complete genomes with minimal assembly fragmentation (Figure 2E). `CheckM2` quality assessment indicated that SymGs had a median completeness higher than 90% and low contamination (typically <0.2%), comparable to the quality metrics observed for genomes in the overall catalog (Supplemental Figure S2B). Many symbiont MAGs are relatively small, streamlined genomes, which contributed to high assembly contiguity (high N50 values) and low contig counts per genome (often only a few contigs; Figure 2E). These statistics are consistent with the biology of well-studied obligate symbionts, which often possess reduced, gene-dense genomes that could be fully or nearly fully recovered from metagenomes. In summary, the SymGs represent a set of genomes that not only spans diverse taxa but also meets high standards of completeness and purity, ensuring that downstream analyses of symbiont biology rest on a solid genomic foundation.

#### 3.2.3. Ecological and geographic distribution of symbiont genomes

The Symbiont Genomes catalog – SymGs, exhibits broad ecological and geographic coverage, demonstrating that symbiotic microbes are ubiquitous across global environments (Figure 2F). SymGs were recovered from environmental samples spanning all major biome categories and continents/oceans (Figure 2F), highlighting a near-global distribution of these genomes. Notably, symbiont MAGs were disproportionately derived from host-associated and aquatic biomes, which together accounted for the majority of SymG source environments (Figure 2F). Host-associated samples (such as animal microbiomes, insect guts, and plant rhizospheres) and aquatic ecosystems (marine and freshwater water) each contributed large numbers of symbiont genomes. Overall, the global and ecological spread of SymGs parallels the broad sampling of the entire genome dataset (Supplemental Figure S2C), but with a clear emphasis on environments involving host organisms or aquatic settings where symbiotic interactions are expected to be especially common.

To further characterize the environmental contexts of the symbiont genomes, we examined the most common biome sources represented in SymGs (Figure 2G). The top contributing biomes included the deep subsurface of terrestrial and aquatic samples. Each of these biomes contributed hundreds to thousands of symbiont genomes, underlining that symbiotic microbes thrive in a variety of settings but are especially prevalent in association with animal hosts and aquatic ecosystems. On the contrary, and as expected, obligate intracellular microbes were not detected in built environments (Figure 2G). Non-human animal digestive systems showed presence of predicted obligate intracellular symbionts, while host-associated (and not obligate intracellular) lifestyles were the predominant in human digestive systems (Figure 2G) which may represent more facultative or environmentally transmitted symbionts.

Collectively, the construction and characterization of the Symbiont Genomes Catalog (SymGs) reveal a remarkable breadth of microbial symbionts. The SymGs comprise over fourteen thousand genomes spanning myriad taxa and including high-quality representatives of known and novel symbiont lineages. By contrasting the SymGs with the broader genome catalog, we observe that symbiotic lineages make up roughly 15%-23% of the sampled microbial diversity, a substantial and previously underappreciated fraction.

### 3.3. Functional traits that differentiate microbial lifestyles and allow accurate prediction of symbionts

To identify the genomic features that contribute to the accurate prediction of microbial lifestyles, we performed a SHAP (SHapley Additive exPlanations) analysis on the annotated features, which revealed distinct functional signatures that differentiate free-living and symbiotic bacteria (Figure 3A). Features were stratified into four quadrants based on SHAP effect and feature detection via HMMs bit score (Figure 3A), highlighting two key sets of lifestyle markers (Figure 3B). Quadrant 1 (Q1) features, defined by positive SHAP effect and low bit score, were typically absent in symbionts and thus indicative of free-living bacteria. In contrast, Quadrant 2 (Q2) features, with positive SHAP effect and high bit score, were predominantly present in symbionts. Analysis of Gene Ontology (GO) term semantic organization revealed functional distinctions between feature sets (Figure 3B; Supplemental Figure S3). Core biosynthetic pathways, including enzymes for essential amino acid and nucleotide biosynthesis (e.g. methionine, purine, and pyrimidine synthesis), were enriched among Q1 features, contributing positively to free-living predictions. This is consistent with symbionts often lacking complete biosynthetic capabilities.

**Figure 3.**
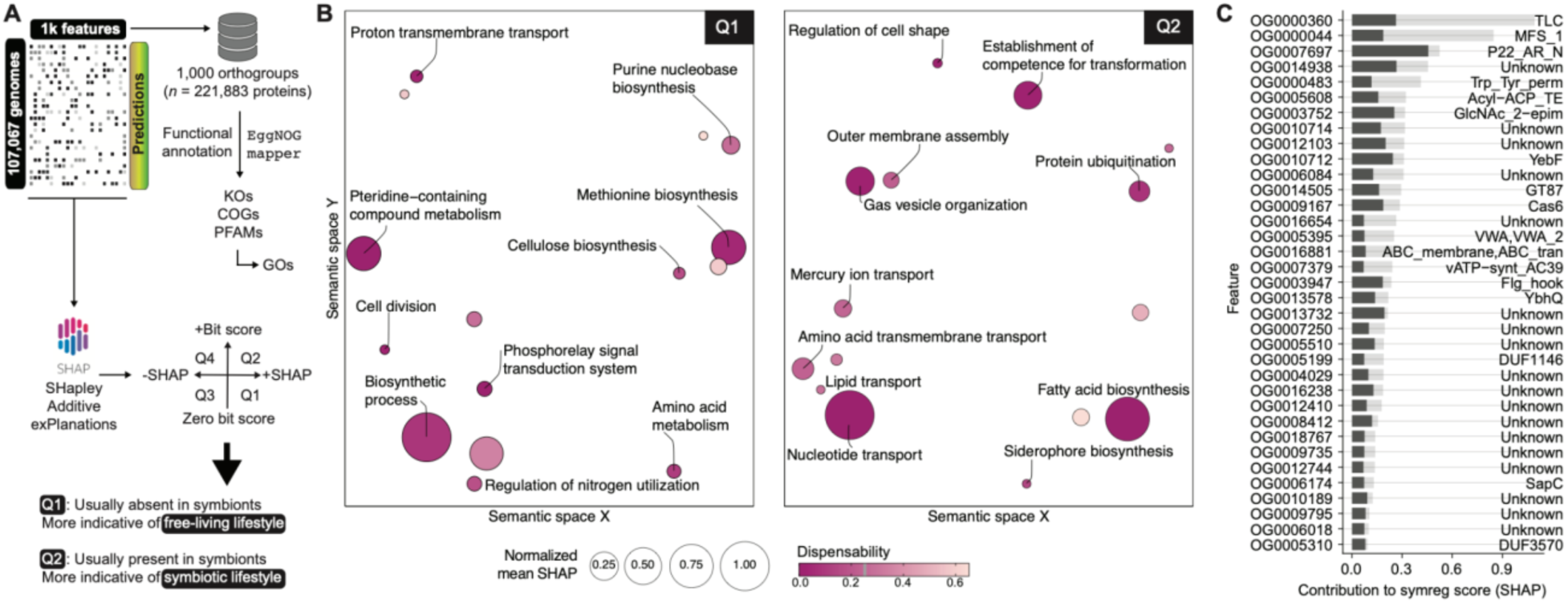
Different functional traits that enable accurate microbial lifestyle predictions by symclatron. (A) Method for functional characterization of features and their contribution to lifestyle prediction. (B) Multidimensional scaling of semantic similarities of the GO terms with highest mean SHAP of features more indicative of free-living (left) and symbiotic (right) lifestyles. Only labels of GO terms with the lowest dispensability (≤ 0.25) were added. Semantic similarities and dispensability of GO terms were computed with REVIGO. (C) Ranked individual contribution of features to the symbiotic lifestyle prediction (only Q2). Mean (solid color bar), and max (translucent color bar) SHAP values of the top 35 features with the highest SHAP mean. PFAM domain annotations are shown on the right.

Conversely, nutrient acquisition and specialized structural functions were enriched among symbiont-associated Q2 features, including numerous nutrient transporters and membrane-associated proteins. Pathways for fatty acid biosynthesis and structural adaptations (e.g. outer membrane assembly) were overrepresented in this set, highlighting adaptations to a host-dependent lifestyle. The top 30 PFAMs most strongly associated with symbiont classification (Figure 3C) were dominated by transport systems and cell envelope biogenesis factors. The differential distribution of core metabolic versus nutrient-scavenging pathways was observed (Figure 3B; Supplemental Figure S3). Importantly, the high SHAP contributions of these biologically coherent traits demonstrate that the model’s accurate lifestyle predictions are grounded in genuine functional differences. Thus, the presence of complete biosynthetic modules or, alternatively, specialized transport and structural systems is a strong determinant of a microbe’s lifestyle.

### 3.4. symclatron allows identification of bacterial and archaeal clades dominated by symbionts

#### 3.4.1. Widespread distribution of symbionts across Bacteria and Archaea

Microbial symbionts are phylogenetically widespread, appearing in numerous branches of the bacterial (Figure 4) and archaeal (Figure 5) trees of life. Nearly all major phyla include some host-associated members, indicating that transitions to symbiosis have occurred repeatedly in diverse lineages. However, the prevalence of symbionts varies greatly among clades, with certain groups especially enriched for symbiotic lifestyles (Figure 4 and 5). For example, the Candidate Phyla Radiation of bacteria (e.g. Patescibacteria) and the DPANN superphylum of archaea (e.g. Nanoarchaeota, Aenigmatarchaeota) consist almost entirely of host-dependent organisms. By contrast, in large phyla like Pseudomonadota or Actinomycetota, symbionts represent a smaller subset of the total diversity, yet even in these groups, multiple subclades (for instance, the order Rickettsiales in Alphaproteobacteria or family Mycobacteriaceae in Actinobacteria) have evolved host-associated lifestyles (Figure 4).

**Figure 4.**
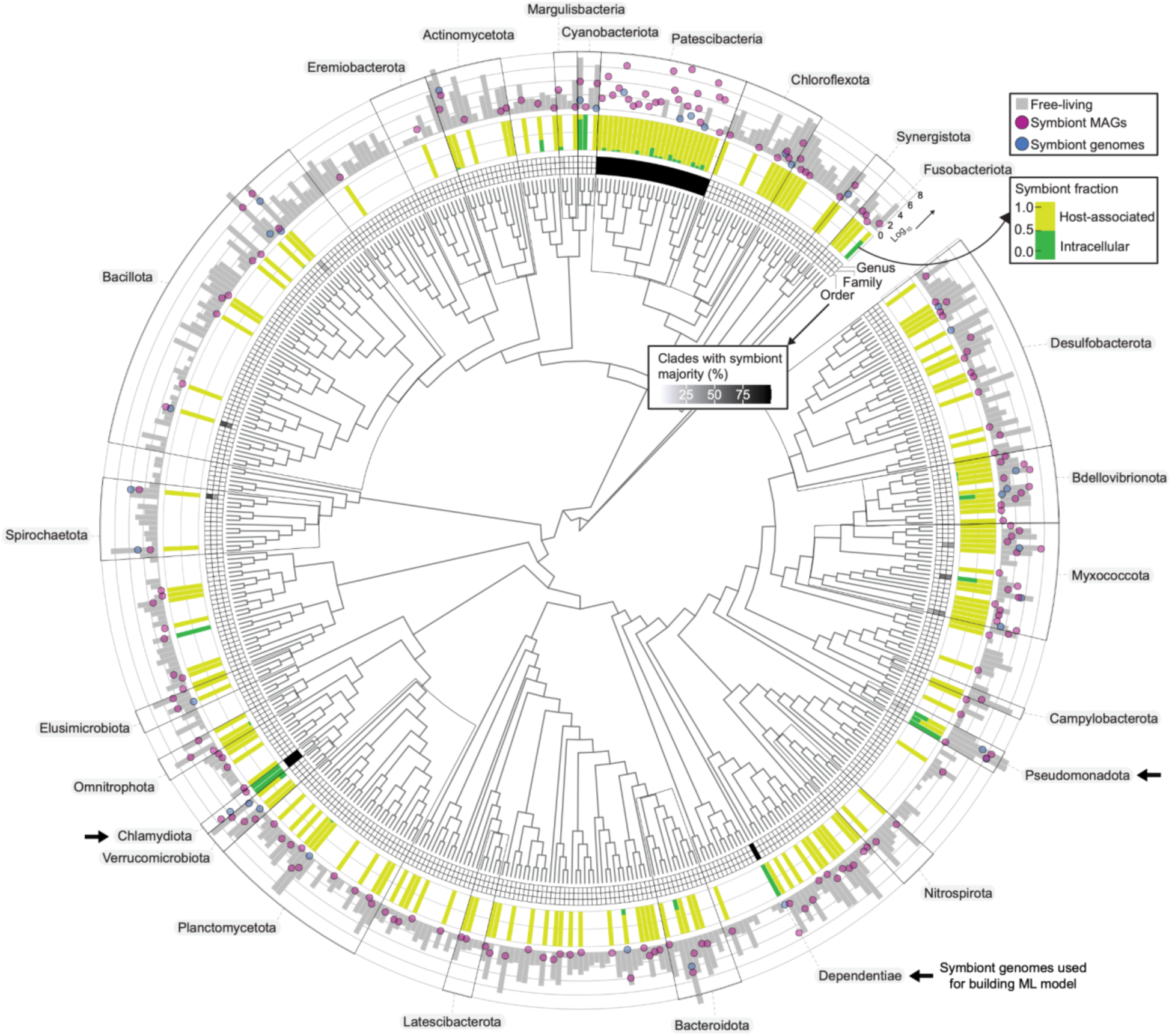
Phylogenetic distribution of the bacterial symbionts. The tree encompasses a representative genome from each class spanning various phyla. The tree was rooted using Fusobacteriota. From the innermost to the outermost rings the following information is displayed: (i) The heatmap ring shows the percentage of symbiont-exclusive clades per class found at the order, family, and genus levels. (ii) The bars represent the specific lifestyle of the symbiont fraction within the class. (iii) The filled dots and the gray bars show the *log_10_*-scaled total number of genomes analyzed in each class. (iv) Clades from which genomes were used in building the classifier and regressor models are designated by black arrows. Additional details on genomes of predicted symbionts and symbiont-exclusive clades are provided in Supplemental Tables S2 and S3.

**Figure 5.**
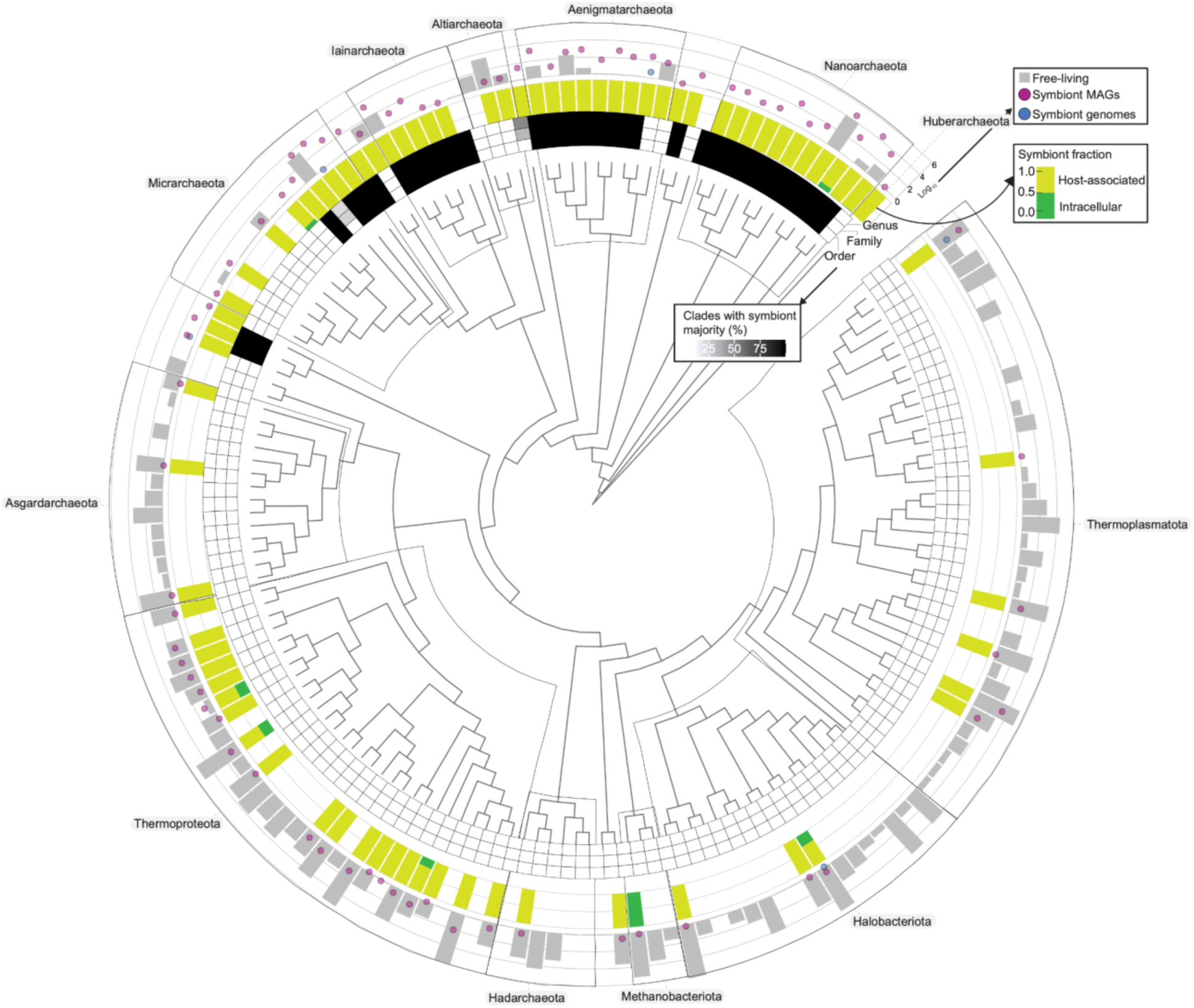
Phylogenetic distribution of the archaeal symbionts. The tree encompasses a representative genome from each order spanning various phyla. The tree was rooted using bacterial outgroups which have been removed from the visualization. From the innermost to the outermost rings the following information is displayed: (i) The heatmap ring shows the percentage of symbiont-exclusive clades per order found at the order, family, and genus levels. (ii) The bars represent the specific lifestyle of the symbiont fraction within the order. (iii) The filled dots and the gray bars show the *log_10_*-scaled total number of genomes analyzed in each order. Additional details on genomes of predicted symbionts and symbiont-exclusive clades are provided in Supplemental Tables S2 and S3.

Our analysis also shows that many symbiont-exclusive clades are dominated by intracellular symbionts (See Supplemental Table S3 for the detailed metrics of every clade analyzed in this study). For instance, the phylum Chlamydiota (Figure 4) is comprised entirely of obligate intracellular bacteria (most are expected to be pathogens of animals based on the current understanding of this clade), and the Dependentiae (formerly TM6) are similarly almost exclusively intracellular symbionts (most are expected to be endosymbionts of protists based on the current understanding of this clade). In other clades, the symbionts are predominantly host-associated but not obligate intracellular, as in the case of certain bacterial clades within the Bacteroidota and Bacillota (formerly known as Firmicutes) phyla that are traditionally known to colonize the gut of animal hosts without residing inside host cells. In sum, the broad phylogenetic survey of symbiont genomes shows that symbiotic relationships have evolved in diverse taxa of the microbial world, with some lineages undergoing especially pronounced shifts to host-dependent, often intracellular, existence in Bacteria (Figure 4) and Archaea (Figure 5).

#### 3.4.2. Symbiont-exclusive clades

Using the above phylogenomic framework, we identified numerous “symbiont-exclusive” clades, defined as taxonomic groups in which ≥90% of the member genomes are predicted to be symbionts with high-confidence (Figure 6A). As expected due to microbial diversification, the majority of symbiont-exclusive clades are found at the lower taxonomic ranks: over a thousand genera consist almost entirely of symbionts. At progressively higher ranks, fewer clades meet the ≥90% symbiont criterion. We detected on the order of tens of classes and hundreds of orders and families that are symbiont-dominated, but only a handful of phyla (Figure 6A). The few top-level groups with pervasive symbionts include well-known obligate symbiont lineages such as Chlamydiota (phylum of obligate intracellular bacteria), as well as several candidate phyla in bacteria and archaea that lack free-living representatives. This pattern indicates that completely host-dependent groups exist at all taxonomic scales, but broad, phylum-level specializations to symbiosis are rare. More commonly, symbiosis evolves as a specialized subset within larger lineages. The prevalence of so many symbiont-exclusive genera and families suggests multiple independent acquisitions of a host-associated lifestyle within each major bacterial phylum. Furthermore, many of these clades have only recently been recognized through metagenomic sequencing (Figure 6A), implying that our knowledge of microbial diversity has expanded to include previously undiscovered symbiont lineages.

**Figure 6.**
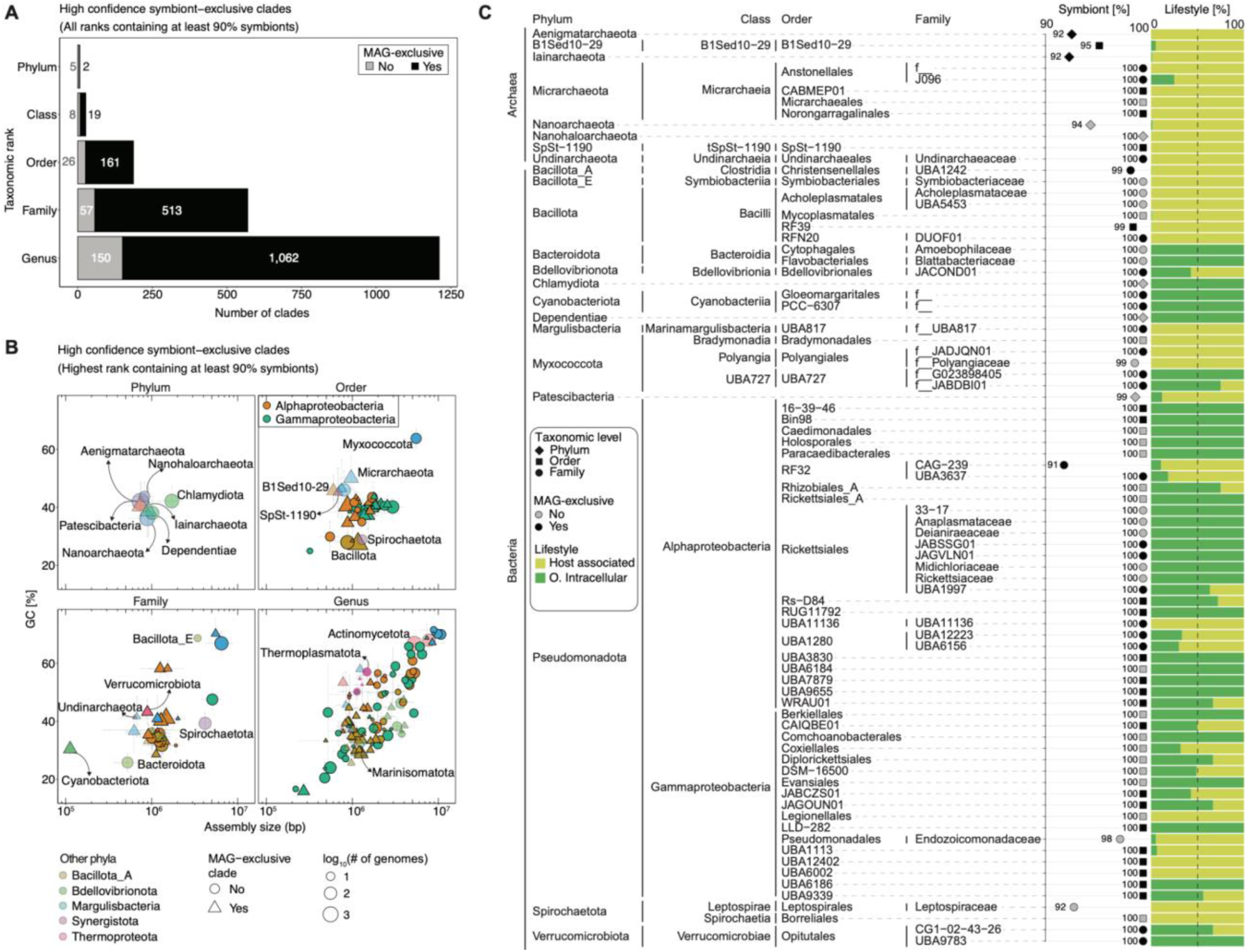
Genomic characterization of symbiont-exclusive clades. (A) Distribution of symbiont-exclusive clades across different taxonomic ranks. A clade was depicted as “symbiont-exclusive” at any taxonomic rank if ≥ 90% of genomes were classified as symbionts with mean confidence of the predictions of the clade being ≥ 0.725 and a standard deviation ≤ 0.2. A clade is depicted as “MAG- exclusive” if it was not previously represented by reference isolates. (B) GC content and assembly size of symbiont-exclusive clades across different taxonomic ranks, from phylum to genus. Dot color indicates the phylum in which we found symbiont-exclusive clades; only Alphaproteobacteria and Gammaproteobacteria are shown at the class level. Each point represents a clade. (C) Lifestyle composition of symbiont-exclusive clades down to the family taxonomic rank. Unknown taxonomic classifications are depicted by the first letter of the taxonomic rank and two underscore signs. The two rightmost columns indicate the fraction of genomes in that clade classified as symbionts, and its percentage distribution of symbiotic lifestyles.

Indeed, a significant fraction of our high-confidence symbiont clades appear to be MAG-exclusive (known solely from metagenome-assembled genomes), highlighting how cultivation-independent approaches are revealing symbionts that were missed in cultivation-based surveys (Figure 6A). Genome characteristics of many symbiont-exclusive clades reflect reductive evolution, but some have larger genomes and high GC content (Figure 6B). There is a pronounced trend toward genome miniaturization: many obligate symbionts in our dataset have genome sizes well under 1.5 Mbp, and some approach the theoretical lower size limit for cellular life (on the order of only a few hundred kilobases). These reduced genomes also tend to have biased nucleotide compositions, often <40% GC. For instance, several known insect endosymbiont clades (Figure 6B) harbor genomes around ∼0.2-0.7 Mb with extreme AT bias (e.g. one lineage at ∼13-20% GC). Such values are in line with the most extreme cases previously reported in endosymbiotic bacteria like *Candidatus* Zinderia insecticola with a 208 kb, 13.5% GC genome (McCutcheon and Moran 2010).

To examine the nature of these symbiont-dominated clades more concretely, we broke down a selection of high-confidence symbiont-exclusive clades (down to family level for clarity), detailing their taxonomy, symbiont percentage, and lifestyle composition (Figure 6C). All of the highlighted clades have at least 90% of members identified as symbiotic, with many at a full 100%. The proportion of host-associated vs. intracellular symbionts is indicated for each clade, illustrating different degrees of integration with host organisms. Some clades are composed almost entirely of obligate intracellular symbionts. For example, the family Blattabacteriaceae (order Flavobacteriales) is 100% symbiotic, consisting of obligate intracellular (traditionally known to be endosymbionts of cockroaches) bacteria that live within host fat-body cells (Vicente et al. 2018). Similarly, the Amoebophilaceae (order Cytophagales) are intracellular symbionts traditionally known to live inside amoeba hosts (Selberherr et al. 2022). In these groups, virtually all symbionts inhabit host cells, indicating an extreme specialization and dependency on the host environment.

In contrast, other clades, while still exclusively host-associated, predominantly contain non-obligate intracellular symbionts (Figure 6C). An example is the order Acholeplasmatales (class Bacilli [former Mollicutes], phyla Bacillota), which includes mycoplasma-like bacteria found on animal and plant hosts; these symbionts require a host to survive but typically reside on cell surfaces or in host fluids rather than intracellularly invading host cells. As a result, Acholeplasmatales genomes, though reduced, are not as minimal as those of strictly intracellular bacteria (Bacillota dots in Figure 6B). Likewise, certain gut bacterial clades like Christensenellales (former Christensenellaceae) within the class Clostridia and phyla Bacillota_A (former Firmicutes) are known to be almost entirely restricted to animal hosts yet remain outside host cells (Waters and Ley 2019) as predicted by us (Figure 6B).

Among the Pseudomonadota symbionts that were used for building the classifier, surprisingly, only Rickettsiales did not meet the criteria for being considered a symbiont-exclusive clade despite its known occurrence as symbionts of a wide variety of eukaryotic species (Figure 6C). Both orders Holosporales and Legionellales did form symbiont-exclusive clades (Figure 6C).

Within Rickettsiales, only Rickettsiaceae, Midichloriaceae, JAGVLN01, JABSSG01, Deianiraeaceae, Anaplasmataceae, and 33−17, formed symbiont-exclusive families of obligate intracellular symbionts (Figure 6C). In contrast, some members of the Rickettsiales family UBA1997 were classified as host-associated. Alternatively, these could also be ectosymbiotic, a lifestyle that has been reported in the Rickettsiales for “*Candidatus* Deianiraea vastatrix” (Castelli et al. 2019).

Overall, our data indicate that most clades dominated by symbionts have gravitated toward intracellular niches (Figure 6C), with puzzling cases like the MAG-exclusive unknown families of Gloeomargaritales and PCC−6307 within Cyanobacteriota (Figure 6C). Nevertheless, in other clades, a significant subset maintain a less intimate, non-obligate intracellular association with their hosts. This variation suggests that while host dependency is the common thread uniting these clades, the mode of symbiosis (intracellular endosymbiosis versus host-association) can differ, likely influenced by the specific ecological context and evolutionary history of each lineage.

### 3.5. Differential metabolic module completeness found in symbiont-exclusive clades compared to their free-living relatives

#### 3.5.1. Widespread loss of biosynthetic pathways in symbiont genomes

We calculated metabolic module completeness for each clade and employed bootstrap resampling to detect major differences between symbiont and free-living groups. 100 random subsamples of genomes were used to generate confidence intervals for each module’s completeness in symbiont vs. free-living cohorts. This analysis ensured that observed differences (e.g., loss of amino acid synthesis or enrichment of nitrogen metabolism) are not due to random genome-content variation but reflect true adaptive divergences.

We observed that, across diverse lineages, many core metabolic pathways are incomplete or entirely absent in symbiont-exclusive clades (Figure 7). For example, symbionts in the families CG1-02-43-26 and UBA9783 within the order Opitutales (phylum Verrucomicrobiota), and the order Borreliales within the phylum Spirochaetota, show pervasive loss of amino acid biosynthesis pathways that are present in related free-living bacteria. Modules for the production of branched-chain amino acids (valine, isoleucine and leucine), as well as other essential amino acids like ornithine/arginine and histidine, are largely missing or pseudogenized in the symbionts (Figure 7). This trend is echoed in the symbionts-exclusive order Borreliales within the phylum Spirochaetota, which lack key enzymes for serine, threonine, and arginine biosynthesis that their free-living relatives retain (Figure 7). We generally observed that symbionts show pervasive loss amino acid biosynthesis pathways that are present in related free-living bacteria. Similarly, symbionts rely on salvage pathways for nucleotides: we found de novo purine and pyrimidine biosynthesis modules (e.g., M00048 and M00051) to be partially or wholly absent in symbiont clades, whereas free-living strains encode these pathways completely (Figure 7).

**Figure 7.**
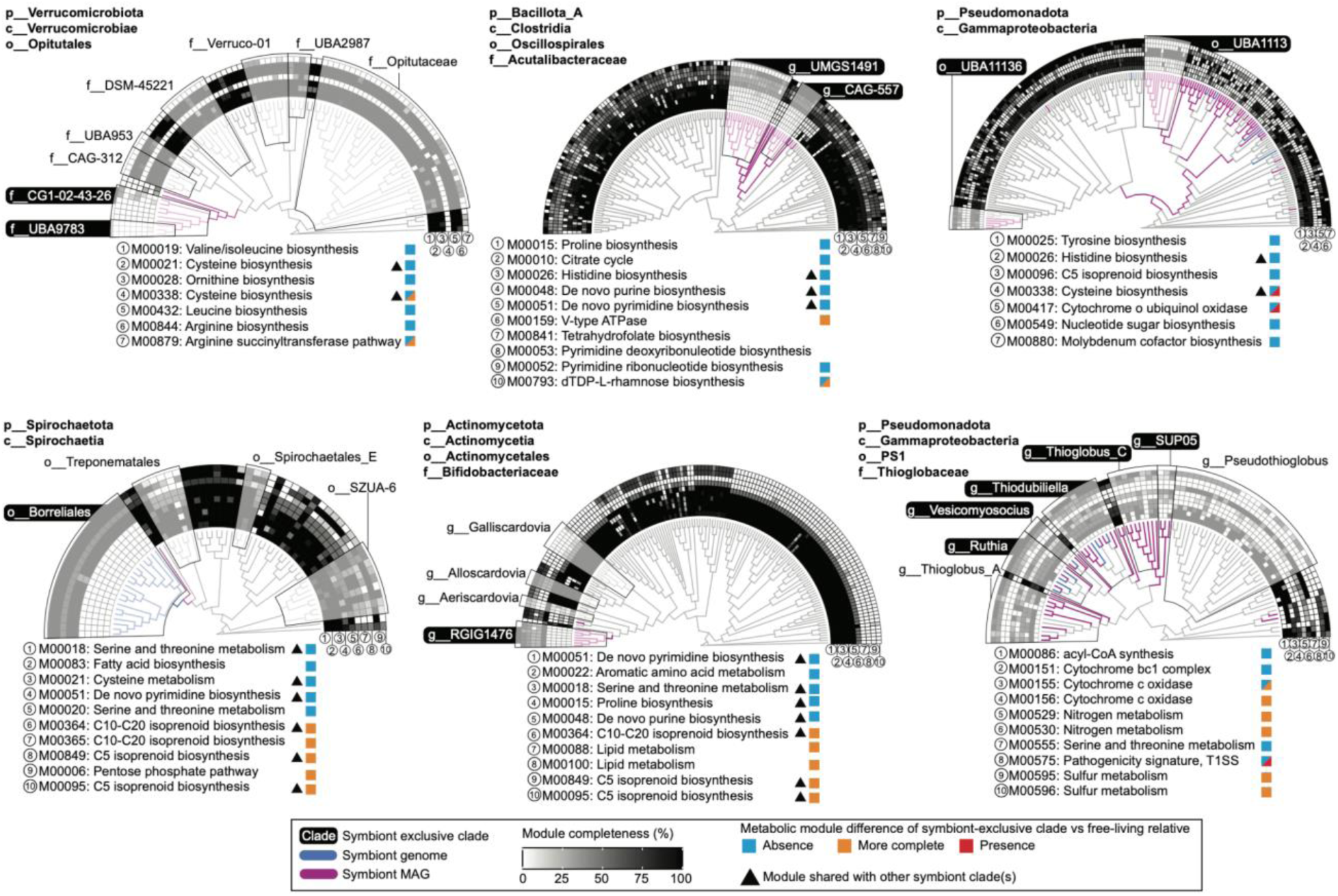
Differential metabolic module completeness between genomes of symbionts and their free-living sister clades. Phylogenomics trees showing clades identified as symbiont- exclusive alongside related taxa. Key metabolic modules were automatically selected based on the bootstrapping approach detailed in Figure S4A. Modules that are shared between two clades are denoted by triangles. Module identification is based on the KEGG database. Each ring of cells around the tree represents a KEGG metabolic module, with fill color indicating the completeness or presence/absence of that module in each genome.

Interestingly, the two symbiont genera UMGS1491 and CAG-557 within the family Acutalibacteraceae (class Clostridia and phylum Bacillota_A) were characterized by having also shed parts of central metabolic pathways like the citrate cycle (TCA cycle), being absent in all examined symbionts within these two clades (Figure 7). Their absent citrate cycle indicates a reduced capacity for aerobic respiration in these symbionts. Nevertheless, they do show a more complete module for V-type ATPase metabolism.

In a contrasting example of related symbiont clades with potentially different hosts, we observed that the symbiont-exclusive order UBA1113 within Gammaproteobacteria retained a complete module cytochrome o ubiquinol oxidase, but the same module was not consistently detected in the related symbiont-exclusive order UBA11136, and was completely undetectable in free-living counterparts (Figure 7). This differential presence of the cytochrome o ubiquinol oxidase module indicates that UBA1113 are still occupying niches with oxygen flux while UBA11136 are most likely adapting to oxygen-poor host environments.

Our data show that the loss of energy-intensive biosynthetic functions indicates that symbionts frequently obtain amino acids, nucleotides, cofactors, vitamins, and other key metabolites (Supplemental Figure S4B) from their hosts (or co-symbionts), a hallmark of metabolic dependency in host-associated bacteria (Boscaro et al. 2017; Nechitaylo et al. 2021; Osvatic et al. 2023). However, some symbiont-exclusive clades can also show more complete modules for these biosynthetic functions (Supplemental Figure S4B), indicating that the selective pressure of the specific host/niche environment is a differential factor in the evolution of metabolic capabilities of symbionts, and reduction of metabolic function is not the only adaptation mechanism of host-associated and intracellular microbial symbionts.

#### 3.5.2. Retention of niche-specific metabolism in symbionts genomes

We observe that despite overall metabolic contraction, symbionts commonly retain or even enrich certain pathways that may confer a selective advantage within the host environment. Intriguingly, some biosynthetic modules are more complete in symbionts than in related free-living taxa (Figure 7). For example, the modules for C5 and C10-C20 isoprenoid biosynthesis, and pentose phosphate pathway in the symbiotic order Borreliales within Spirochaetota are consistently more complete.

Nitrogen-metabolism modules in symbiont-exclusive genera within the family Thioglobaceae (class Gammaproteobacteria) are more complete than in free-living relatives (Figure 7), indicating potentially nitrogen-poor and sulfur-poor diets of their hosts or the symbionts’ retention in nitrogen acquisition modules likely alleviates their hosts’ dietary deficiencies, providing a source of usable nitrogen. Likewise, we found that sulfur metabolic modules are retained in these symbiont clades, probably reflecting adaptation to anoxic, sulfate-rich gut environments. Most likely, the nitrogen and sulfur chemoautotrophic metabolism that is fully present, enables the host to exploit inorganic energy sources in harsh environments like deep-sea vents. In the symbiotic genera CAG-557 within the class Clostridia (Figure 7), the dTDP-L-rhamnose biosynthesis module, often critical for the virulence of human pathogens (Mistou, Sutcliffe, and van Sorge 2016), is consistently complete in contrast to other symbiont clades and free-living relatives.

Overall, we observe that symbionts thereby may allow hosts to live in habitats or utilize resources that would otherwise be inaccessible, illustrating an adaptive advantage of maintaining certain metabolic modules. In general, the genes that remain in reduced symbiont genomes (Supplemental Figure S4B) tend to be those directly involved in metabolic exchanges with the host, like nutrient provisioning or waste recycling, highlighting the tight metabolic interdependence of host and symbiont.

## 4. DISCUSSION

Our study leverages large-scale genome-resolved metagenomics and machine learning to explore the global diversity of microbial symbionts of eukaryotes. By compiling and analyzing a comprehensive catalog of genomes primarily composed of metagenome-assembled genomes (MAGs), here we show the extensive diversity and pervasiveness of bacterial and archaeal symbionts across Earth’s biomes, an idea long hypothesized (Margulis 2008; Husnik et al. 2021), but not documented until now.

We developed the `symclatron` software designed for classification of genomes. One of the main criteria for `symclatron` development and optimization was to minimize the false positive rate for symbionts prediction, which provides a conservative trustworthy ground for the host-associated and obligate intracellular predictions made here. Nevertheless, `symclatron` showed remarkable accuracy by correctly classifying symbiotic lifestyles in clades such as Bipolaricaulota and Omnitrophota, consistent with recent literature (Tandon et al. 2022; Seymour et al. 2023). Another example includes “*Candidatus* Azoamicus ciliaticola” (GTDB: f UBA6186, o UBA6186), a Gammaproteobacterial endosymbiont that uses denitrification to generate ATP for its ciliate host (Graf et al. 2021); “*Ca.* Azoamicus ciliaticola” was the first described member of a much larger clade of protist endosymbionts (Speth et al. 2024). In our analysis, we were able to identify 21 genomes from the order UBA6186 that are part of our SymGs catalog. `symclatron` predicted all the symbionts within the UBA6186 order as intracellular (Supplemental Table S3).

`symclatron` correctly predicted several lineages of archaeal symbionts affiliated with DPANN lineages, despite `symclatron` being trained on only bacterial symbionts. One such example is Nanoarchaeota (Wurch et al. 2016; Jarett et al. 2018), which associate obligately with other Archaea within the Thermoproteota. Many members of Thermoproteota were also identified as symbionts by `symclatron`, however, our classifier predicted the known Thermoproteota hosts *Ignicoccus hospitalis* (Paper et al. 2007) and Acd1 (Podar et al. 2013) to be correctly free-living. Further, it correctly identified the symbiotic and free-living lifestyle of the Huberarchaeota-Altiarchaeota symbiosis, respectively (Schwank et al. 2019). Finally, it predicted symbiont-exclusive clades for Micrarchaeota which were recently described as an example for Archaea-Archaea interactions (Krause et al. 2022). `symclatron`’s capability in predicting prokaryote-prokaryote associations is potentially supported further by the high prevalence of predicted symbionts in deep subsurface samples. This biome is known to harbor eukaryotic organisms such as fungi and protists, but typically in very low abundance (Borgonie et al. 2015; Bhattacharjee et al. 2023). The substantial proportion of predicted symbionts becomes more plausible when considering the likelihood of prokaryote-prokaryote associations in this environment.

We detected host-associated clades, as in the case of the MAG-exclusive Bipolaricaulota (Figure 4). Although MAGs from this phylum have been recently sequenced from coral hosts (Tandon et al. 2022), it is important to note that they are phylogenetically distant from the clades used for feature extraction in our classifier. In agreement with recent results (Seymour et al. 2023), our classifier also predicted both host-associated and obligate intracellular clades within the Omnitrophota. Within Patescibacteria (Rinke et al. 2013); also referred to as Candidate Phyla Radiation (Brown et al. 2015), our classifier predicted most (99% genomes) as symbionts. This suggests that, despite their limited metabolic capacities and likely dependency, some members of this phylum might not require other organisms to thrive, which would support recent findings (Beam et al. 2020; Chaudhari et al. 2021).

Due to the capability of microbial symbionts to be a key driver of complex life evolution by influencing the adaptive radiation of host lineages to a variety of environments, understanding symbiont diversity and pervasiveness is essential (McCutcheon 2021). Some symbiosis, especially the transition from free-living to obligate intracellular lifestyles, leads to an irreversible genome reduction and metabolic streamlining of the symbiont genome (N. A. Moran 1996; Wernegreen 2005; Toft and Andersson 2010; McCutcheon and Moran 2011; Bennett and Moran 2015). However, although it is clear symbiont genomes are reduced during coevolution with their host, there is evidence that gene gain may enable the initial symbiotic interaction that leads to prolonged association. For example, a recent discovery indicated gene gain (Dharamshi et al. 2023) as an evolutionary mechanism in the transition to symbiosis for Chlamydiota. Our study also reveals a streamlining of certain biochemical networks in symbiont genomes, such as central metabolic pathways and biosynthesis of nucleotides and amino acids, presumably sustained by hosts. Yet, we also report here differential function presence in symbiont-exclusive clades compared to their free-living relatives across several pathways and from multiple clades (Figure 7). One clear example is the increased completeness found in the metabolic module M00417: Cytochrome o ubiquinol oxidase (part of the bacterial electron transport chain complex IV), in the symbiont-exclusive order UBA1113 within Alphaproteobacteria, linking our results to recent findings of gain of function by Chlamydiota in their transition to endosymbiotic lifestyle (Dharamshi et al. 2023). These findings were further extended through the detection of presence or more complete metabolic modules, in several symbiont-exclusive clades, involved in pathways related to cofactor, vitamins, and ATP metabolisms.

The reported differential presence of metabolic capacity challenges the traditional paradigm of genome streamlining in host-associated and intracellular bacteria, hence we provide extensive evidence of evolution towards selective retention of metabolic capabilities in symbionts. These findings extend our understanding of the adaptive outcomes that genomes of symbionts underwent (or are currently undergoing) to sustain ecological complexity (Muñoz-Gómez, Kreutz, and Hess 2021) and thrive in their specific hosts and their niches. Different mechanisms of gene transfer have been recently described in bacterial symbionts, such as gene transfer agents in endosymbionts within the Alphaproteobacteria (George et al. 2022), ancestral vertical inheritance accompanied by gene innovations in Rickettsiales (Schön et al. 2022), and gene gain in Chlamydiota (Dharamshi et al. 2023). The evolutionary mechanisms of metabolic retention, and potential gain of functions, found in symbiont lineages in our collection remain to be investigated.

We collectively demonstrate that microbial symbionts have evolved streamlined genomes that sacrifice broad metabolic independence in favor of a pared-down set of functions tailored to the host environment. Key biosynthetic pathways have been lost in multiple symbiont lineages, indicating a convergent evolutionary response to the host-provided niche. Conversely, symbionts preferentially retain or enhance pathways that fulfill host needs or exploit host habitat resources, underscoring a fine-tuned metabolic complementarity between host and symbiont (Figure 7 and Supplemental Figure S4). These findings highlight how intimate symbiotic associations drive metabolic specialization and co-dependency, ultimately enabling hosts to occupy ecological niches that would otherwise be unattainable.

In conclusion, our findings fill a major gap in the understanding of the bacterial and archaeal symbionts, and offer new global perspectives on the ecological, evolutionary, and biochemical implications of microbial symbiosis, and is expected to lay the groundwork for future studies aimed at exploring the novel functional roles and mechanisms of symbionts in understudied hosts from various difficult-to-access ecosystems.

### 4.1. Limitations of the study and future directions

Our study provides novel insights into the microbial symbionts of eukaryotes on a global scale, but it is important to acknowledge certain limitations and outline directions for future research. One of the limitations lies in the inherent biases of metagenomic sampling. Despite our extensive dataset there may have been an underrepresentation of certain hosts, geographical locations, or biomes. Furthermore, challenges linked to accessing extreme environments, and economical and sociopolitical limitations likely contribute to uneven distribution of sampling locations. Future efforts could focus on targeted sampling after host enrichment and from underrepresented biomes to provide a more balanced view of the microbial symbionts.

While `symclatron` represents a powerful tool in the context of culture-independent microbiology, it is grounded on existing genomic data and annotations, which may not capture the full spectrum of microbial symbiosis. Metagenomics poses a challenge as well, in particular as short-read sequencing yields often only fragmented assemblies with partial genome recovery, potentially obscuring the complete picture of symbiotic capabilities. Further, MAGs and SAGs may contain contaminant contigs missed by state-of-the art contamination screening tools, or contain chimeric sequences, which may result in a potential bias in gene content recovery (Chang et al. 2024). The application of long-read sequencing technologies and improved assembly algorithms could enhance genome completeness and the accuracy of lifestyle predictions.

Other limitations can be linked to our training data set for which we selected experimentally verified symbionts with sufficient genome coverage, biasing our data set toward a set of symbionts of eukaryotes. However, even though we did not train `symclatron` on symbioses between prokaryotes due to the lack of literature on such interactions and available genomes, `symclatron` still predicted several recently described cases of symbioses among Bacteria and/or Archaea. Future versions of `symclatron` will systematically incorporate these and other newly described symbioses as part of the training data.

Another challenge is that functional interpretation of symbiotic diversity is based on the current understanding of metabolic pathways and their ecological roles. As our knowledge expands, re-evaluation of these pathways in the context of symbiosis will be necessary. In particular, investigation of expression and regulation of these pathways *in situ* would offer deeper insights into the functional dynamics of symbiotic interactions. Lastly, the evolutionary aspects of symbiosis revealed by our study invite further exploration into the genetic mechanisms that drive the transition from free-living to symbiotic lifestyles. Integrating phylogenomics, systems biology, and ecological data, could elucidate the mechanisms, evolutionary pressures, and historical contingencies that shape symbiotic associations.

## 5. METHODS

### 5.1. Definition of symbionts, host-associated, and obligate intracellular

In this study, we adopted the definitions recently delineated by Husnik et al. (2021) to characterize symbiotic relationships. We define a symbiont as an organism that maintains a spatially close and temporally prolonged association with a host organism of a different species. Symbionts can be broadly categorized as host-associated or, more specifically, as obligate intracellular. Host-associated symbionts encompass both facultative endosymbionts, which can replicate inside a host cell but also independently of a host, and ectosymbionts (or epibionts), which inhabit the exterior of their host, often attached to the host’s surface. In contrast, obligate intracellular symbionts represent a subset of host-associated symbionts that thrive and replicate exclusively inside a host cell. To better reflect the continuum of symbiosis and the blurred boundaries of these definitions, we use the term “intracellular” instead of ℌobligate endosymbiont” throughout the manuscript. While a prediction of “host-associated” could also indicate that a microbe can thrive inside or attached to a eukaryotic host cell facultatively.

In our training data, we assigned higher scores to known obligate intracellular symbionts (label = 2, “obligate intracellular”) and lower scores to known facultative endosymbionts and ectosymbionts (label = 1, “host-associated”).

Syntrophic relationships, where two organisms cooperate to metabolize a substance that neither could break down alone, fall outside of this definition of symbionts. Syntrophs, while metabolically interdependent, are not typically considered symbionts in the strict biological sense because in contrast to symbionts that engage in intimate, often obligate interactions with a host, syntrophic microbes cooperate primarily through environmental metabolite exchange without physical integration or long-term coadaptation (Sieber, McInerney, and Gunsalus 2012). High-impact theoretical frameworks emphasize that true symbiosis involves specific host-microbe associations maintained across generations and shaped by selection for mutual benefit (Bright and Bulgheresi 2010; López-García, Eme, and Moreira 2017). Broader ecological interactions like syntrophy, despite conferring functional advantages, are better conceptualized as community-level cooperation rather than symbiosis (Douglas and Werren 2016; Nancy A. Moran and Sloan 2015).

### 5.2. AI/ML-based classification of microbial lifestyles

We developed a machine learning classifier to systematically classify microbial genomes based on their lifestyles. First, all the proteins (*n* = 1,642,823) found in known genomes of symbionts within the clades detailed in Figure 1A were subjected to `OrthoFinder` (v2.5.4) (Emms and Kelly 2019) to detect orthogroups of proteins across the different microbial clades of symbionts (Figure 1A). Only protein sequences from orthogroups with at least 5 sequences were retained. Sequence alignment and trimming were performed using `MAFFT` (v7.481) and `trimAl` (v1.4), respectively, to prepare for the construction of Hidden Markov Models (HMMs). A total of 20,063 HMMs were generated to represent the protein orthogroups (Figure 1A).

To construct a database of labeled genomes according to lifestyles (Figure 1B), we downloaded one (or two when available) genomes from each bacterial genus in the IMG/M (accessed in October 2022). The genomes (*n* = 6,751, Supplemental Table S1) were then labeled (free-living: 5,959; host-associated: 409, intracellular: 383) by scrutinizing for lifestyle categories using manual literature searches.

Next, every labeled genome was artificially fragmented to emulate various levels of genome completeness (50%, 60%, 70%, 80%, and 90%) and fragmentation lengths (10 kb, 30 kb, and 50 kb) with `shred.sh` from `BBMap` (v39.01) (Bushnell 2014). Gene prediction on these fragmented genomes was conducted using gene-calling with `Prodigal` (v2.6.3), to identify proteins after the fragmentation iterations. The predicted proteins from labeled genomes were then searched against the 20,063 HMMs generated from the symbiont orthogroups using `hmmsearch` to obtain bitscores. In order to keep consistent and reproducible results, all `hmmsearch` runs were with the parameters: “`-E 1000`, `--incE 1000`, and `-Z 61295632` to guarantee the entire set of results were output independently of potential E-value biases generated by data base size. This parameter was chosen intentionally with the goal of maximizing sensitivity. We aimed to capture all potential homologs, including those with only remote similarity, ensuring that potentially relevant but divergent features were not missed.

While a high E-value threshold allows many hits, the statistical significance and strength of each hit are still captured by the HMMER bitscore. These E-values parameters were chosen to maximize sensitivity and capture even remotely homologs, allowing the machine learning step to interpret these features in context and produce reliable predictions. Finally, the 20,063 HMMs deriving from the symbiont genomes were then utilized to search against the proteins of labeled genomes (Figure 1B) via `hmmsearch` (v3.3.2) (Eddy 2011).

For the AI/ML component (Figure 1D), we constructed a training matrix with 20,063 features (from the HMMs), encapsulating the bitscores of every genome and its simulated genomes from the labeled data set. These features were fed into two gradient-boosted decision tree models: a classifier (named `symcla` for symbiont classifier) and a regressor (named `symreg` for symbiont regressor), using the GPU version of `xgboost` (v1.7.4) (T. Chen and Guestrin 2016) for `Python`. For computational efficiency, we used computing nodes with NVIDIA A100 (40GB) GPUs on the Perlmutter supercomputer of the National Energy Research Scientific Computing Center (NERSC) (https://www.nersc.gov/, US Department of Energy – Office of Science). The regressor

The two models predict whether a genome falls within the symbiosis continuum (Fig 1C), ranging from free-living to host-associated, then obligate intracellular lifestyles, labeled as 0, 1, and 2, respectively. On one hand, the symbiont classifier (`symcla`) predicts the lifestyle according to the probability score of each class. While, on the other hand, the symbiont regressor (`symreg`) assigns a regression score based on the continuous scale in the range from 0.1 to 2.0.The models are designed to predict the positioning of a given genome along a symbiosis continuum, as illustrated in Figure 1C. This continuum encompasses lifestyles ranging from free-living to host-associated, culminating in obligate intracellular lifestyles, which are categorized numerically as 0, 1, and 2, respectively. The symbiont classifier (symcla) operates by assigning a probability score to each class, thereby predicting the lifestyle. Conversely, the symbiont regressor (symreg) employs a continuous scale, assigning a regression score within the range of 0.1 to 2.0.

The neural network- based classifier was then built by first extracting a set of numeric features from each genome and its classifications by the classifier and the regressor (e.g., predicted scores from regression and classification models, completeness, and distance to training data measures). These features were then combined into a feature matrix, and the corresponding labels were one- hot- encoded. The dataset was split into training and validation subsets at an 80:20 ratio, and a StandardScaler was fitted on the training data to normalize each feature’s distribution. This step helped stabilize the subsequent model training process. Next, a feed- forward neural network architecture was defined. It consisted of an input layer feeding into three hidden layers with Rectified Linear Unit (ReLU) activation functions, each followed by BatchNormalization (to stabilize gradients) and Dropout (to reduce overfitting). The output layer was a three- neuron softmax layer, producing probability estimates for the three lifestyle classes. To address potential class imbalance, class weights were calculated and applied during training. The model was optimized using Adam (learning rate = 0.001) with categorical crossentropy loss, and training was monitored with early stopping based on validation loss.

The final softmax probability from the neural network serves as an initial “confidence score” for each prediction, which can be used directly or thresholded to balance accuracy and coverage. The final confidence score was derived by first extracting the probability value of the predicted lifestyle class (the class with the highest probability from the neural network output). This base confidence was then adjusted by two biological context factors: predictions for taxonomically distant proteomes (weighted minimum Euclidean distance >80) received a 0.8 penalty factor. The final confidence score was calculated by multiplying the base confidence by the penalty factor when applicable, resulting in a metric ranging from 0 to 1 that integrated both statistical certainty and biological context. The final AI/ML framework composed of the classifier, the regressor, the extraction of genome features, and the neural network was named `symclatron`.

The culmination of the AI/ML framework efforts was the development of a novel software tool, `symclatron` (https://github.com/NeLLi-team/symclatron). In the `symclatron` software, we developed and included one module for rapid estimation of genome completeness through the detection of UNI56 marker genes as a close, but computationally efficient, approximation to CheckM2 completeness estimates (Supplemental Figure S5).

### 5.3. Building a non-redundant genome catalog of Earth’s microbiomes and reference genomes

Metagenomic bins were obtained via metagenome binning and gene calling on assembled contigs from 31,152 public environmental sequencing data projects through the IMG/M system up to April 2023. Only contigs with length equal to or greater than 5 kbp were processed.

Metagenome samples without associated depth data were removed. Binning was performed on each sample with `MetaBAT2` (parameters: --minContig 5000 --minClsSize 20000 --cvExt) (Kang et al. 2019), yielding 5,635,662 bins recovered from metagenomes, cell enrichments and single amplified genomes (SAGs) (Figure 2A) from which 136,784 passed the quality criteria of `CheckM2` (v1.0.2) (Chklovski et al. 2023) estimates for completeness equal or higher than 50% and contamination lower than 5%. Gene calling was performed on each bin with Prodigal (v2.6.3) with the `-p meta` parameter (Hyatt et al. 2010).

Adding to our catalog, reference genomes and MAGs available through publicly available repositories were downloaded from the IMG isolates repository (*n* = 115,969) (I.-M. A. Chen et al. 2023) and from the bacterial and archaeal representative datasets of GTDB (*n* = 85,205) (Parks et al. 2022). All genomes and MAGs were subsequently subjected to `CheckM` (v1.1.3) with the UNI56 markers for detection of of universal markers required for further phylogenomics reconstruction, `CheckM2` (v1.0.2) for quality assessment, and `seqkit` (v2.5.1) (Shen et al. 2016) for assembly characterization. All genomes and bins with less than 50% UNI56 markers or `CheckM2` completeness lower than 50% or `CheckM2` contamination higher than 5% were removed from the genome catalog.

Cluster representatives were determined based on the highest assembly quality and dereplication through `skani` (v0.1.4, parameters: `ANI` ≥ 95% `align_fraction_ref` ≥ 50% and `align_fraction_query` ≥ 50%) (Shaw and Yu 2023). A single metric for assembly quality “*Q*” was computed as: *`Q`* `=(0.5*completeness_checkm2)+[0.2*(100- contamination_checkm2)]+(0.3*completeness_UNI56)`to quantitatively represent the quality criteria applied. Then the cluster representatives were selected as the ones with the highest *Q*, and if ties were found, the assemblies with the highest number of genes predicted and the highest N50 were selected. The de-replicated high quality genome catalog was finally composed of 107,067 genomes and bins; these were further taxonomically characterized with `GTDB-Tk` (v2.4.0) (Chaumeil et al. 2022) from the GTDB Release 220.

### 5.4. Systematic cross-validation and calibration of final predictions

In order to ensure that our predictive model generalizes well across diverse microbial clades, we meticulously cross-validated predictions accounting for taxonomic diversity of the labeled genomes using the classifier and the regressor. We adopted a stringent strategy where clades were entirely omitted from the training dataset to evaluate each model’s performance on unseen data. Each of the training and testing matrices includes an iterative process of removing entire clades corresponding to different taxonomic levels, ranging from phylum to species levels (*n* = 11,384, one per clade found). For example, when benchmarking at the phylum level, we remove the entire set of genomes belonging to the phylum Pseudomonadota from the training data set, we trained on all other phyla, and used all Pseudomonadota only in the testing data set; the same was applied for all clades at every taxonomic rank. This validation technique ensures that each model’s predictive power is not artificially inflated by over-represented taxonomic groups. Each training iteration of the classifier used a learning rate of 0.1 and 1,000 estimators. The aggregated outputs of each of the 11,384 cross-validation exercises were used as input to the neural network for final classification calibration and confidence estimation (Figure 1E).

### 5.5. Feature contribution to microbial lifestyle prediction

The feature contributions to the model’s predictions were analyzed utilizing the Shapley Additive exPlanations (SHAP) methodology, implemented within the `shap Python` library. SHAP values, which provide a breakdown of the impact of each feature on the model’s output, were computed exclusively for the `symreg` predictions. This decision was made because the inherent structure of regression models allows for a systematic and interpretable analysis of how individual features contribute to either increasing or decreasing the predicted regression score. This approach unveils the impact of individual genomic features on the predictive model’s output, providing an interpretable mechanistic basis for the classification of microbial lifestyles by decomposing the `symreg` prediction into the sum of effects from each feature, allowing us to quantify the impact of individual genomic attributes. We computed the SHAP values for each of the 1,000 features of each one of the predicted symbionts. Next, all proteins in the 1,000 most relevant features (*n* = 221,883 proteins) were functionally annotated using `EggNOG-mapper` (v2.1.10) (Cantalapiedra et al. 2021), effectively assigning each feature to KEGG KOs (KEGG ortholog groups) (Kanehisa and Goto 2000) COGs (clusters of orthologous groups) (Tatusov et al. 2000) and PFAMs (Protein FAMilies) (Finn et al. 2014), allowing for the biological interpretation of the features and aligning the machine learning insights with known biological functions. Gene Ontology (GO) terms were mapped from PFAM using `pfam2go` (https://current.geneontology.org/ontology/external2go/pfam2go). Our framework employs a quadrant-based approach to interpret the contributions of genomic features towards the classification of microbial symbiotic lifestyles. As shown in Figures 3A, this framework delineates features into four quadrants, each representing a unique combination of presence or absence and impact on the model’s predictive score:

Quadrant 1 (Q1): Features that contribute positively to the `symreg` score mainly through absence (`hmmsearch` bitscore < 20). More indicative of free-living lifestyles.

Quadrant 2 (Q2): Features that contribute positively to the `symreg` score mainly through presence (`hmmsearch` bitscore ≥ 20). More indicative of symbiotic lifestyles.

Quadrant 3 (Q3): Features that contribute negatively to the `symreg` score mainly through absence (`hmmsearch` bitscore < 20). Indicative of symbiotic lifestyles.

Quadrant 4 (Q4): Features that contribute negatively to the `symreg` score mainly through presence (`hmmsearch` bitscore ≥ 20). Indicative of free-living lifestyles.

The bitscore threshold of ≥ 20, used to determine the potential presence of a given domain within a protein sequence, was established based on the distribution of Gathering threshold for sequence and domain used by the InterPro PFAM HMM profiles (e.g. https://www.ebi.ac.uk/interpro/entry/pfam/PF00010/logo/).

### 5.6. Phylogenomics of the bacterial and archaeal symbionts

We inferred phylogenomic trees using `NSGTree` (https://github.com/NeLLi-team/nsgtree) from concatenated protein sequences from at least half of the UNI56 universal marker genes. In `NSGTree`, UNI56 markers are identified with `hmmsearch` (v.3.3.2, parameters: `--cut_ga`). Next, we removed genomes if less than 50% of the markers were absent, or if at least one marker was present in more than 4 copies, or if more than 30% of the markers were present in multiple copies. Sequences were aligned using `MAFFT` (v.7.310) and trimmed with `trimAL` (v.1.4.1) using a mild trimming to minimize the loss of potentially informative sites (parameter: `- gt 0.1`). Concatenated protein sequences were then used to reconstruct a species tree using `IQ-TREE` (v.2.0.3) with the LG+C60+R8+F model. Trees were visualized with the `ggtree R` package (Yu 2020).

### 5.7. Definition of symbiont-exclusive clades

We designated a clade as symbiont-exclusive if at least 90% of its members were finally classified as symbionts, the clade included a minimum of three genomes, and the mean confidence of the clade was at least 0.725 with standard deviation of the confidence below 0.2. This designation was made using the `symclatron` final classification. We considered clades at all taxonomic levels, as determined by GTDB-Tk (Chaumeil et al. 2022).

When symbiont-exclusive clades were compared against their free-living relatives (“sister clades”), as in the case of the metabolic module completeness analyses, free-living clades were only considered if they included at least 5 genomes and if clade members, at the same or higher taxonomic levels, had predominantly free-living lifestyle predictions.

### 5.8. Comparative analysis of metabolic module completeness

We assessed the completeness of metabolic modules of symbiont-exclusive clades and their free-living relatives, based on the metabolic module definitions of KEGG (Kanehisa and Goto 2000). The KEGG KOs functional annotations of every genome were extracted from the results of `EggNOG-mapper` (v2.1.10) (Cantalapiedra et al. 2021); when more than one KO was assigned to a protein, all the KOs were unnested and considered individual KOs for downstream analyses. The list of KOs per genome was then passed to the script `ko_mapper.py` from `MicrobeAnnotator` (v2.0.5) (Ruiz-Perez, Conrad, and Konstantinidis 2021) to map KOs to their respective modules and compute metabolic module completeness per genome.

We compared the completeness of KEGG metabolic modules between predicted symbiont- exclusive clades (i.e., clades with ≥ 90% symbionts and ≥ 3 members) and their predicted “free-living” sister lineages by firstly collecting all available free- living relatives at the same phylogenetic rank (the “sister” clade). If there were at least 5 free- living members, 100 bootstrap samples of the same metabolic module were drawn for the symbiont and the free-living groups. Then, for each metabolic module, the average completeness of the bootstrap samples from symbionts is then compared against the average completeness of the bootstrap samples from free- living relatives, yielding a distribution of completeness differences. From this distribution, mean Δ (delta) completeness and 95% confidence intervals are computed for each module.

## 6. SUPPLEMENTAL INFORMATION

Supplemental information (figures and tables) has been submitted together with this manuscript and will be publicly available immediately upon publication.

The symclatron software, and all the accompanying models and data bases, are available at https://github.com/NeLLi-team/symclatron.

All FASTA files of genomes that are part of the SymGs catalog will be hosted at https://portal.nersc.gov/cfs/nelli/symgs/ and will be publicly available immediately upon publication.

All the environmental DNA sequencing samples from the IMG/M system, in which we found symbionts, are individually cited in Supplemental Table S4. Data providers were individually contacted according to the JGI’s Data Policy at least one month before submission of the manuscript.

## 7. ACKNOWLEDGMENTS

We would like to thank the expert advice of Jean-Marie Volland regarding the symbiosis continuum and Antonio Camargo on machine learning benchmarking. The work conducted by the U.S. Department of Energy Joint Genome Institute (https://ror.org/04xm1d337), a DOE Office of Science User Facility, is supported by the Office of Science of the U.S.

## 8. AUTHOR CONTRIBUTIONS

Conceptualization, F.S. and J.C.V.; software J.C.V., F.S., and E.W.W.; investigation, J.C.V., F.S., Y.M.V., and S.Q.; writing – original draft, J.C.V., Y.M.V, and F.S.; writing – review & editing all authors; visualization, J.C.V., F.S., and Y.M.V.; funding acquisition, F.S. and T.W.; supervision, F.S.; project administration, F.S.

## 9. DECLARATION OF INTERESTS

The authors declare no competing interests.

---

**Supplemental Figure 1.**
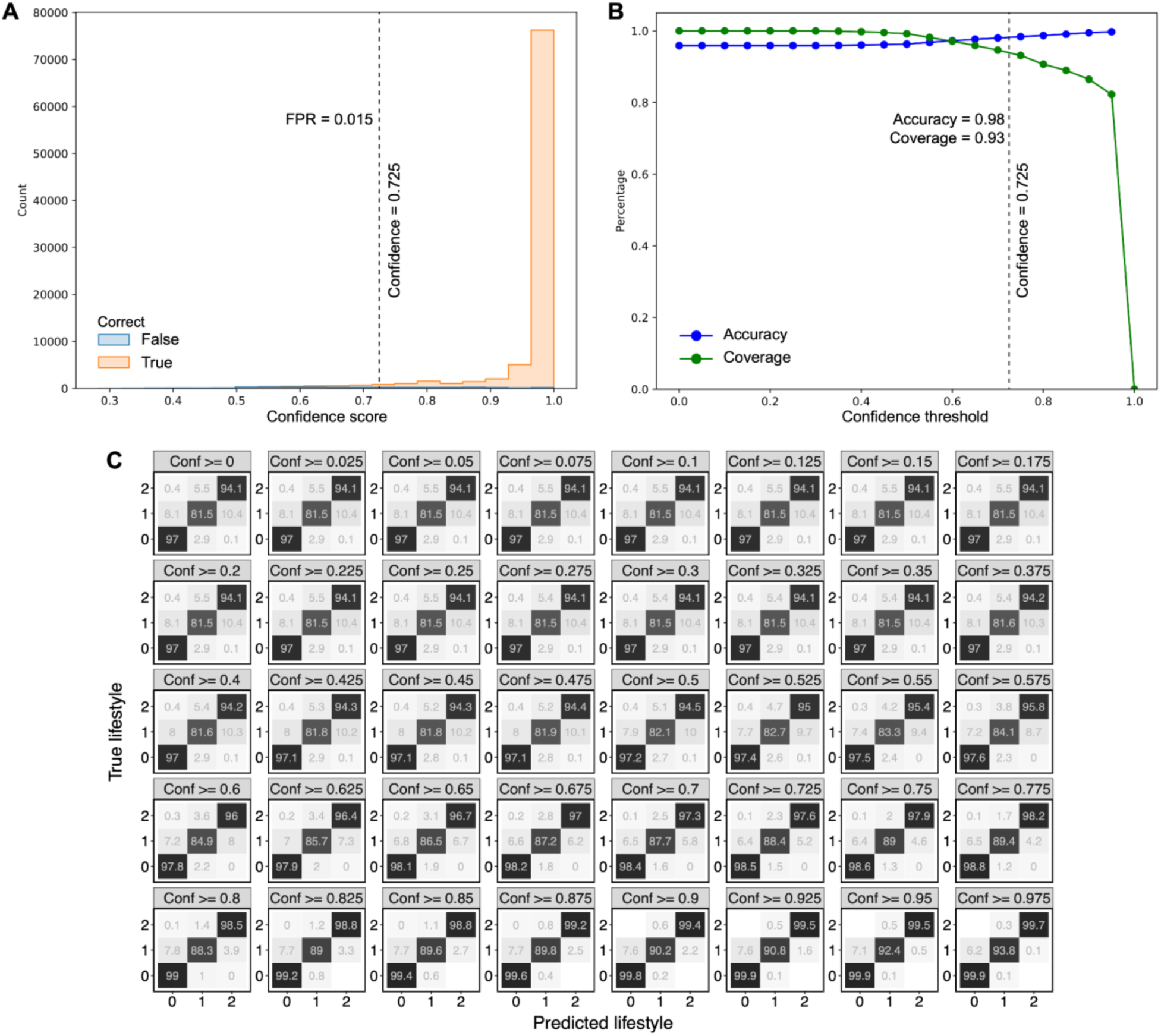
Performance metrics of the lifestyle prediction model. (A) Histogram of the model’s confidence scores for all predictions, colored by whether each prediction was correct (orange) or incorrect (blue). The vertical dashed line marks the chosen confidence threshold of 0.725, which corresponds to a false positive rate (FPR) of 0.015. (B) Classification accuracy (blue) and coverage (green) as functions of the confidence threshold. The vertical dashed line at 0.725 indicates the operating point where accuracy is 0.98 and coverage is still above 0.9 (exactly 0.93). (C) Confusion matrices for the three lifestyle classes (0 = free-living, 1 = host-associated, 2 = obligate intracellular) at progressively stricter minimum confidence cutoffs (in 0.025 increments). The data shown corresponds to the calibration across all taxonomic ranks.

**Supplemental Figure 2.**
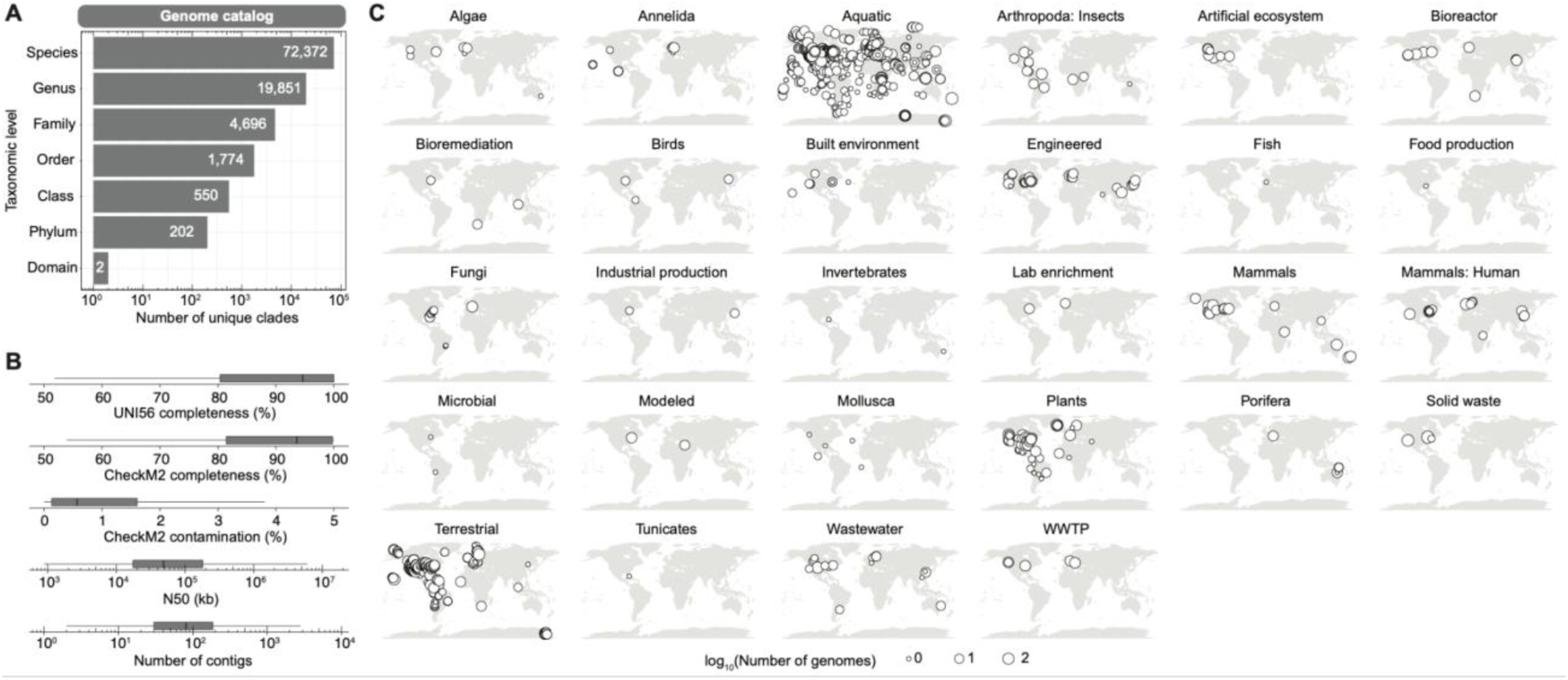
Overview of the complete genome catalog and its global distribution. (A) Taxonomic breakdown of the clades identified in the entire genome catalog. (B) Quality assessment metrics of the entire genome catalog. (C) Global distribution of genomes in the entire genome catalog across different biomes. Geographical locations and environment data is only shown for metagenomes with available metadata in the IMG/M.

**Supplemental Figure 3.**
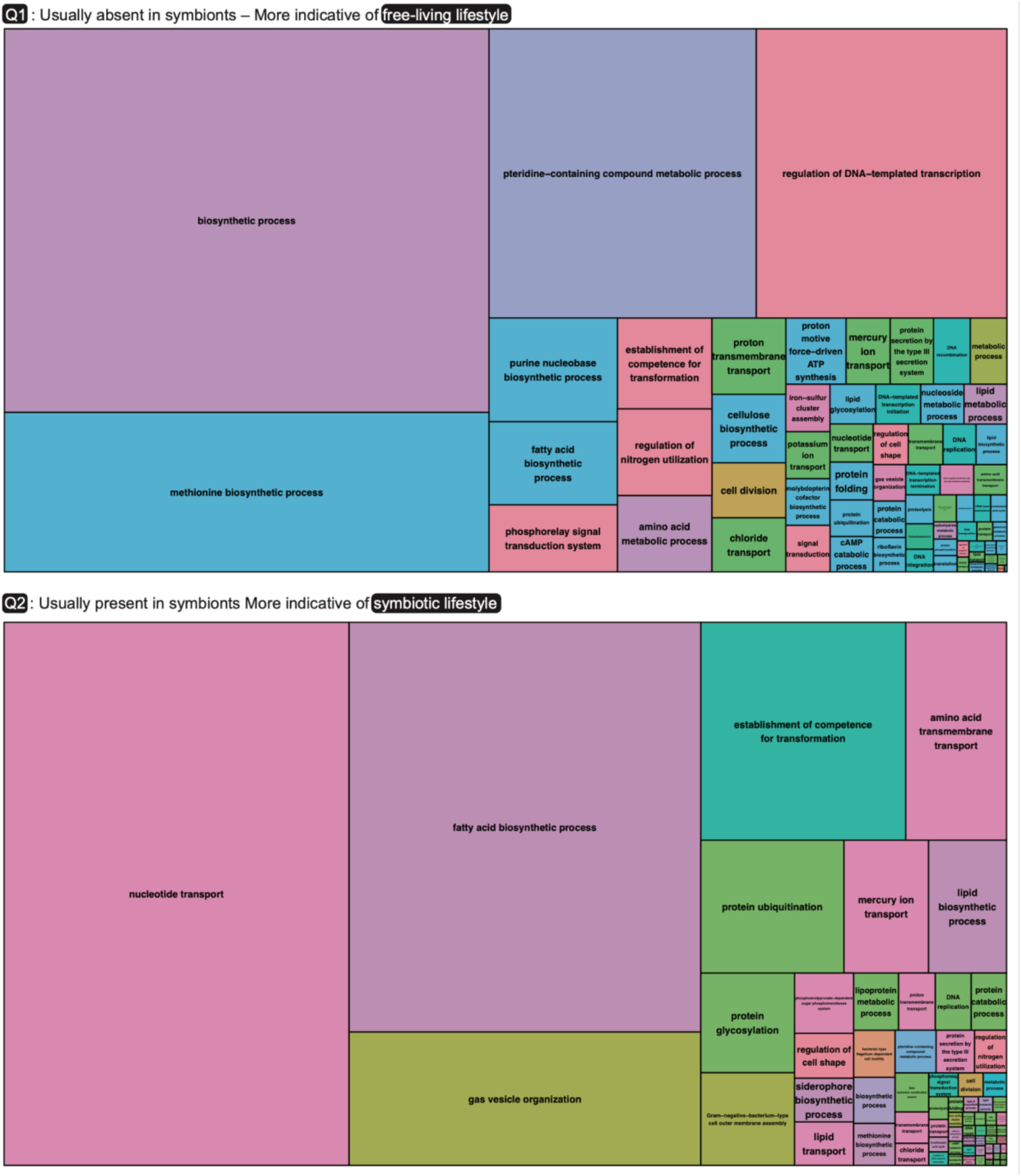
REVIGO treemaps of enriched GO terms for quadrants Q1 and Q2. Each treemap shows representative biological processes after redundancy reduction with REVIGO, grouped by semantic similarity and sized according to the frequency or significance of their respective GO term.

**Supplemental Figure S4.**
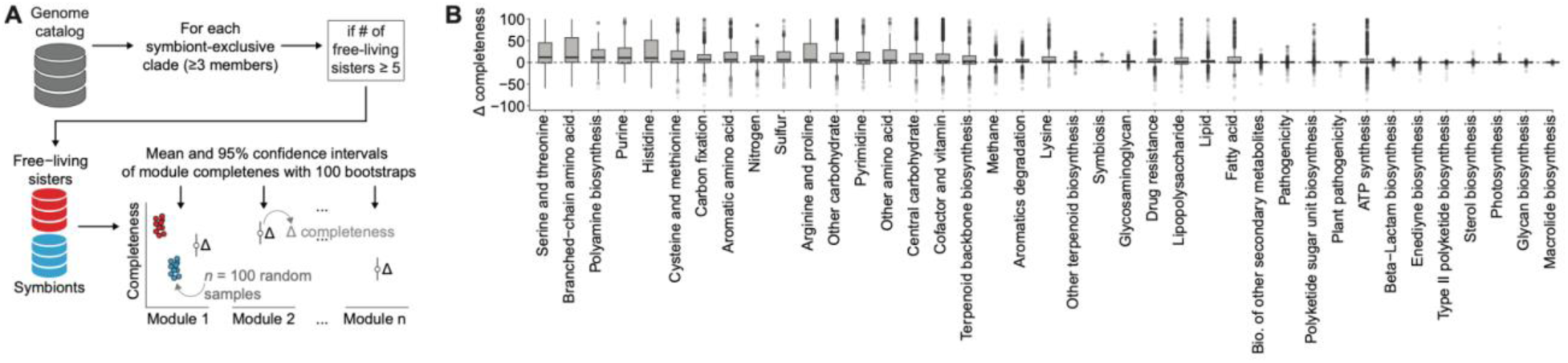
Method developed for automatic detection of differential metabolic module completeness in symbiont- exclusive clades versus their free- living sister lineages. (A) Schematic of the bootstrapping method for detection of differential metabolic module completeness between symbionts and free-living sister clades. For each high-confidence symbiont- exclusive clade containing at least three members, a “sister” set of free- living genomes is identified. If at least three free- living genomes exist in the sister clade, the completeness of each module is compared between symbionts and free- living relatives. One hundred bootstrap samples of free- living genomes are used to estimate the mean difference in module completeness (Δ completeness) with 95% confidence intervals. (B) Differential metabolic module completeness across major metabolic pathways. Each data point corresponds to a symbiont- exclusive clade’s difference from its free- living sister lineage per module, with the horizontal dashed line at 0 indicating no difference. Positive values (above 0) suggest higher module completeness in the free- living set, whereas negative values indicate higher completeness in the symbionts.

**Supplemental Figure S5.**
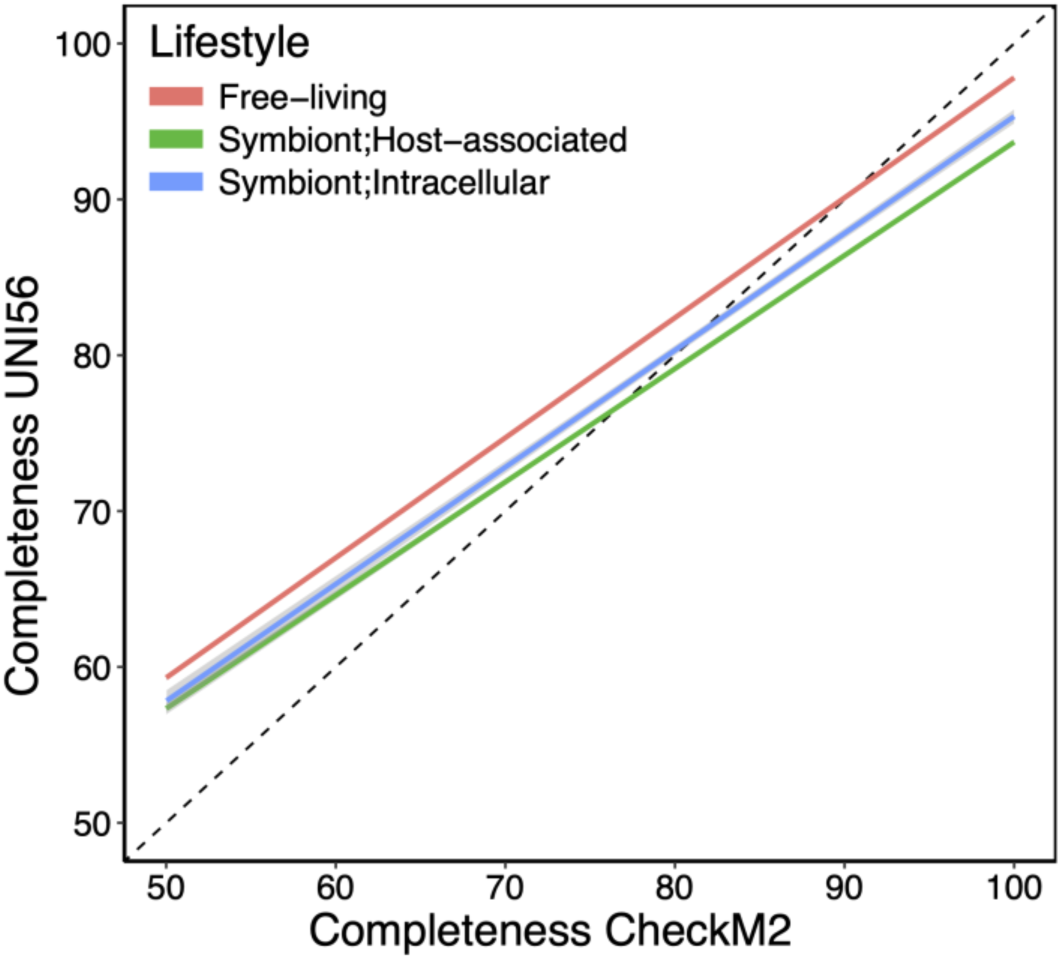
Comparison of genome completeness estimates from CheckM2 and UNI56 markers across different predicted lifestyles. The dashed diagonal line indicates a 1:1 relationship. Fitted trend lines for each lifestyle show the correspondence between the two completeness metrics.

## Notes

https://github.com/NeLLi-team/symclatron

https://portal.nersc.gov/cfs/nelli/symgs/

### Competing Interest Statement

The authors have declared no competing interest.

